# Heterotrimeric Gβ subunit orchestrates flavonoid metabolism governing drought-tolerance in groundnut (*Arachis hypogaea* L.)

**DOI:** 10.64898/2026.09.18.752550

**Authors:** Jismon Jose, Kalyani Prasad, C Vismaya, Swathi Ghantasala, Sudhakar Reddy Palakolanu, Swarup Roy Choudhury

## Abstract

Groundnut (*Arachis hypogaea*) is an important oilseed crop, and its production is severely affected by drought worldwide. Improving the drought-tolerance of groundnut without compromising its agronomic traits is a major challenge for researchers. Here, a meta-transcriptome analysis identified tandemly duplicated Gβ subunits (*AhGβ1* and *AhGβ3*) as early signaling genes under water deficit conditions. Heterotrimeric G-protein β-subunit is a major signaling element that can regulate diverse responses in plants. Heterologous complementation studies in Arabidopsis Gβ mutant (*agb1-2*) demonstrated evolutionary conservation of AhGβ function, while the transgenic overexpression and RNAi knockdown in groundnut established *AhGβ1* as a positive regulator of plant growth, yield, and drought tolerance. Integrated transcriptomic, transcription factor regulatory network, and metabolomic analyses demonstrated that *AhGβ1* coordinates drought-responsive transcriptional and metabolic reprogramming characterized by suppression of cell wall-associated processes and activation of phenylpropanoid, flavonoid, and isoflavonoid biosynthesis. This reprogramming resulted in enhanced accumulation of antioxidant flavonoids, including quercetin and rutin, reduced ROS accumulation, and improved drought tolerance. Furthermore, exogenous application of these flavonoids enhanced drought recovery, highlighting their role as protective downstream metabolites. Collectively, our findings establish *AhGβ1* as a conserved regulator linking early G-protein signaling with transcriptional and metabolic reprogramming to maintain redox homeostasis during drought stress and identify promising molecular and metabolic targets for improving drought resilience in groundnut.

## 1. Introduction

Groundnut (*Arachis hypogaea* L.) is one of the world’s most important oilseed legumes due to its high lipid and protein content. Nearly 70% of global groundnut cultivation occurs in arid and semi-arid regions where water availability is highly unpredictable (Reddy *et al*., 2003). Consequently, drought is one of the most severe environmental constraints limiting groundnut productivity and causes yield loss up to 85% under severe stress (Pokhrel et al., 2025; Reddy et al., 2003). With increasing climate change and frequent occurrence of water scarcity, improving drought tolerance while maintaining agronomic performance has become a major objective for groundnut researchers.

Over the past two decades multiple transgenic efforts have made to improve groundnut drought-tolerance. Overexpression of stress-responsive transcription factors, genes associated with osmotic adjustment, ion homeostasis, and antioxidant defense, has improved drought adaptation in groundnut (Bhatnagar-Mathur et al., 2007; Bhatnagar-Mathur et al., 2014; Bhauso et al., 2014; He et al., 2020; Kiranmai et al., 2018; Li et al., 2021; Pandurangaiah et al., 2014; Patel et al., 2017; Pruthvi et al., 2014; Qin et al., 2013; Qin et al., 2011; Ramu et al., 2016; Rao et al., 2017). However majority of these studies employed heterologous genes previously characterized in other plants. Although numerous transcriptome studies have revealed drought-responsive genes in groundnut (Huang *et al*., 2020; Jiang *et al*., 2021; Wang *et al*., 2021; Zhao *et al*., 2021), the endogenous signaling mechanisms required for adaptation during drought remain largely unknown due to lack of their functional characterization. Therefore understanding these endogenous signaling networks is essential for developing strategies for improving drought resilience in groundnut.

Heterotrimeric G-proteins are major signal transduction complexes that regulate a wide array of developmental and stress-responsive processes, including drought stress in plants. Comprising the Gα, Gβ, and Gγ subunits, these complexes function as molecular switches to transduce extracellular signals into intracellular responses. They are involved in essential processes such as seed germination, organ development, hormone perception, pathogen defense, and tolerance to abiotic stresses (Chakravorty *et al*., 2015; Chen *et al*., 2006; Colaneri *et al*., 2014; Trusov *et al*., 2005; Ullah *et al*., 2002; Ullah *et al*., 2001; Zhang *et al*., 2008). Among these, the Gβ subunit has gained particular attention for its regulatory role, as it interacts with Gγ to form the functional Gβγ dimer that orchestrates multiple signaling outputs. Increasing evidence from model plants highlights the critical role of the heterotrimeric G-protein β subunit (Gβ) in plant responses to drought stress, especially through the modulation of abscisic acid (ABA) signaling, stomatal behavior, reactive oxygen species (ROS) detoxification, and downstream stress-responsive pathways (Fan *et al*., 2008; Jeon *et al*., 2019; Liu *et al*., 2017; Xiong *et al*., 2023; Yu and Assmann, 2018; Yu *et al*., 2018; Zhang *et al*., 2015).

Studies in model plant *Arabidopsis thaliana* have provided the mechanistic framework for understanding Gβ-mediated drought signaling. The single canonical Gβ subunit, AGB1, regulates developmental plasticity as well as ABA-mediated inhibition of stomatal opening, thereby reducing transpirational water loss during drought (Fan et al., 2008; Wang et al., 2001; Wang and Botella, 2022). More recent studies demonstrated that AGB1 interacts with receptor-like kinases such as FERONIA to regulate guard-cell signalling through rapid alkalinization factor (RALF) peptides (Yu *et al*., 2018). In addition, Gβ positively regulates ROS detoxification by enhancing antioxidant enzyme capacity, thereby protecting cellular integrity under water deficit (Liu *et al*., 2017). Functional conservation of Gβ-mediated drought responses has subsequently been reported in wheat, rice, mulberry, tomato and pea, indicating that this signaling module is conserved across plant species (Bhardwaj et al., 2020; Wang and Botella, 2022; Xiong et al., 2023; Zhang et al., 2015). However, whether these mechanistic insights can be translated to legume crops such as groundnut, and how Gβ signaling coordinates downstream transcriptional and metabolic reprogramming during drought adaptation, remain largely unknown.

In the current study, we investigated the biological function of groundnut Gβ subunits in drought adaptation by integrating Arabidopsis complementation, transgenic approaches in groundnut, transcriptomics, metabolomics, physiological characterization, and gene regulatory network analysis. Our objective was to determine whether the conserved functions of Gβ identified in Arabidopsis extend to groundnut and to identify the downstream regulatory processes associated with Gβ-mediated drought adaptation. This study provides a framework for understanding how G-protein signaling contributes to stress adaptation in an important legume crop.

## 2. Results

### 2.1. Meta-transcriptome analysis reveals *AhGβ1* and *AhGβ3* as early drought stress-responsive signaling components in groundnut

We combined and analyzed four different groundnut drought stress RNA-seq datasets, spanning different time points (15) and genotypes (8), using a meta-analysis approach to identify the early-and late-drought stress-responsive genes (Supplementary Figure 1A, Supplementary Table 1). We analyzed early (up to 5 days of drought, 7 comparisons) and late (from 7 days of drought, 8 comparisons) datasets separately. The early time-point comparisons resulted in the maximum number of differentially expressed genes (DEGs) with 2221 up-regulated (early up) and 1805 down-regulated genes (early down) (Figure 1A; Supplementary Table 2). Whereas, the late-time point comparisons resulted in fewer DEGs with 236 up-regulated (late up) and 133 down-regulated genes (late down) (Figure 1A; Supplementary Table 3). Further, the KEGG pathway analysis of this set of genes suggested differential regulation of secondary and glutathione metabolism at early time points and maintenance of these pathways at late time points (Supplementary Figure 1B). Moreover, the downregulated genes from both time points implied the suppression of photosynthesis-related pathways. To understand the major signaling components involved in the early and late responses, we have performed Mercator annotation for the up and down-regulated genes from both time points (Supplementary Figure 1C). This multi-process regulation analysis indicated that most of the processes occurred during the early time points, including calcium, phosphoinositide and G-protein-dependent signaling, programmed cell death (PCD), SnRK1-kinase regulatory system, etc. (Figure 1B). Among these signaling components, the G-protein-dependent signaling is known to be involved in drought stress regulation in model plants. The expression pattern of different G-protein subunits showed that the extra-large Gα protein XLG12 and AhGβ1/3 were consistently up-regulated across early time points, and XLG7, XLG11 and NDL1/2/3 (N-MYC Downregulated-Like) were invariably down-regulated in the early time points (Figure 1C, Supplementary Figure 1D). The genome-wide analysis of the allotetraploid groundnut genome confirmed the presence of multiple heterotrimeric G-protein subunits homologs including 2 Gα, 6 Gβ, 12 Gγ, 12 XLGs and 2 RGS (Supplementary Figure 2A). Further, we focused on the Gβ subunits of groundnut as they are involved in the regulation of both developmental and stress signaling. The relative expression analysis of all six *AhGβ* genes established early drought stress-induced expression of *AhGβ1* and *AhGβ3* (Figure 1D). In addition, treatment with 100 μM ABA, as well as 20% PEG in the seedlings, displayed early transcriptional upregulation of *AhGβ1* and *AhGβ3*, followed by a decline at later time points. (Figure 1E & F). Moreover, all six *AhGβ* subunits showed induced expression during the oxidative stress simulated by 200 mM mannitol (Supplementary Figure 2B).

**Figure 1.**
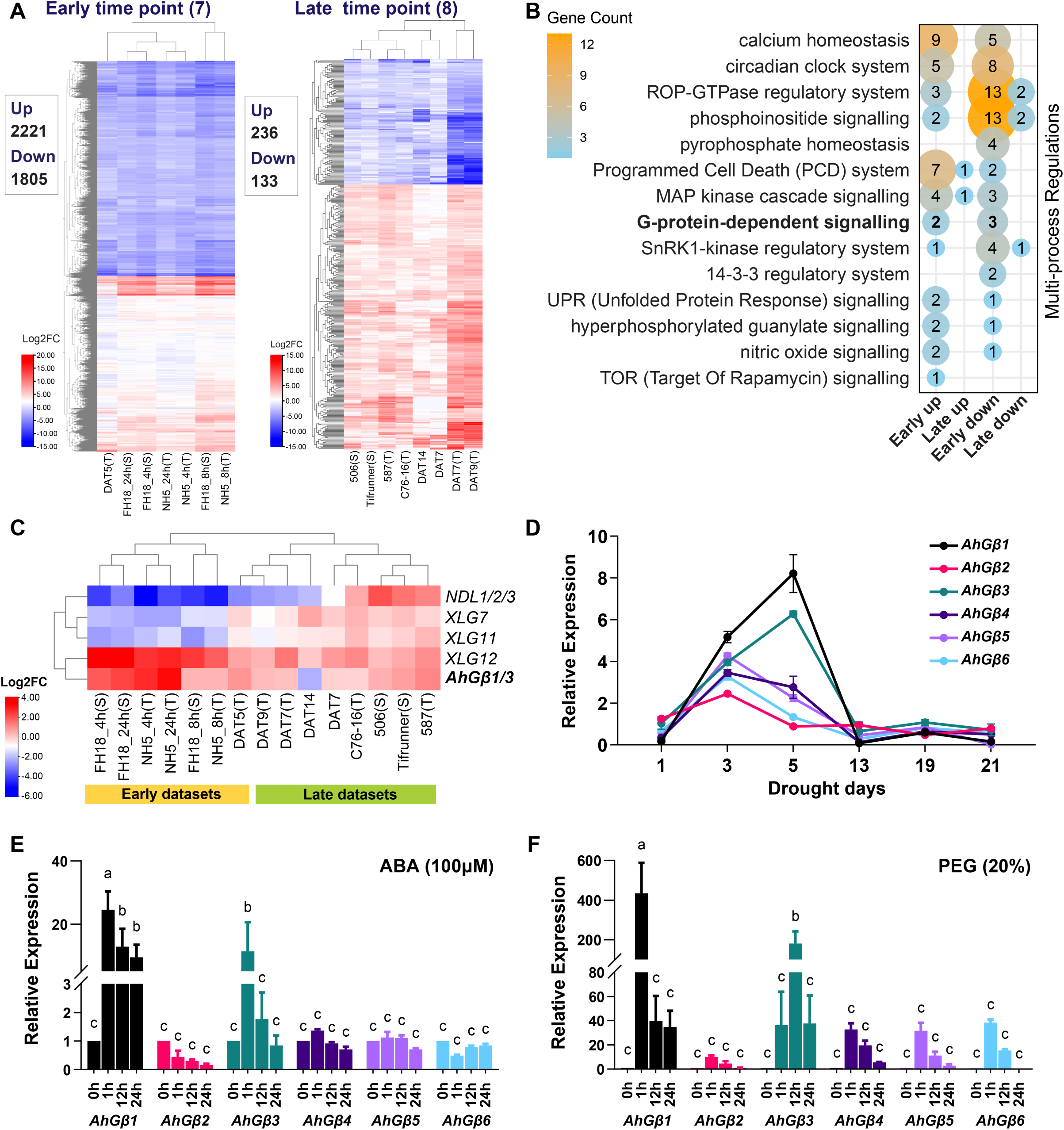
Meta-transcriptome analysis revealed *AhGβ1/3* as an early stress-responsive gene. (A) Heat-maps show the log2FC expression of early- and late-time-point DEGs identified in the meta-analysis. The upregulated and downregulated gene numbers are represented in the left-side boxes, colour scale denoting the log2FC value ranges. (B) The bubble plot exhibits the number of genes in different multi-process regulations across up- and down-regulated genes at early and late time points. Colour scale represents the gene number. (C) The heatmap displays the log2FC expression of major G-protein components identified from meta-analysis across early and late time points; the color scale denotes the log2FC values. (D) The line graph represents the relative expression of all six AhGβ genes across different days of drought stress. (E-F) The bar graphs display the relative expression of all six AhGβ genes at different time points after treatment with 100 pM ABA (E) and 20% PEG (F). The letters on top of bars indicate the statistically significant differences (p<0.05, two-way AN OVA with Fisher’s LSD test).

Further comparative genomic mapping with *A. duranensis* (A genome) and *A. ipaensis* (B genome) indicated that two Gβ orthologs were contributed by both the A- and B-sub genomes of *A. hypogaea* (Supplementary Figure 2C). Evolutionary tracing by conserved syntenic relationship was observed among *AhGβ2*, *AhGβ4*, *AhGβ5* and *AhGβ6* subunits (Supplementary Figure 2D). In contrast, *AhGβ1* and *AhGβ3* showed no detectable syntenic relationship with other Gβ members. Whole-genome duplication (WGD) analysis using MCScanX (multiple-collinearity scan tool kit X) classified *AhGβ1* and *AhGβ3* as tandem or proximal duplicates, whereas all other Gβ subunits were categorized as products of WGD or segmental duplication events (Supplementary Figure 2D). Collectively, these results confirmed that tandem-duplicated *AhGβ1* and *AhGβ3* are the primary stress-responsive subunits in *A. hypogaea*, signifying their potential roles in early abiotic stress signaling. In addition, all six *AhGβ* displayed plasma membrane and nuclear localization similar to their Arabidopsis homolog *AGB1* (Supplementary Figure 3).

### 2.2. *AhGβs* complement stress-sensitive phenotypes of Arabidopsis Gβ mutant *agb1-2*

To elucidate the functional differences among the six *AhGβ* genes, we classified them into three clades; *AhGβ1*/*AhGβ3*, *AhGβ2*/*AhGβ4*, and *AhGβ5*/*AhGβ6* based on the sequence similarity and phylogenetic relationship (Supplementary Figure 2A and 4). One homolog from each clade (*AhGβ1*, *AhGβ4*, and *AhGβ5*) was used in the complementation study in Arabidopsis Gβ mutant *agb1-2* background (Figure 2A). All three *AhGβ* complementation lines exhibited improvement in developmental morphology compared to *agb1-2* mutants. The reduction in rosette diameter observed in *agb1-2* mutant was partially complemented by all three *AhGβs*. Rosette diameter measurements recorded ∼4.6 cm in *agb1-2*, ∼7.5 cm in wild-type (WT) or Col.0 and between ∼6.0 to 7.0 cm in the *AhGβ1*, *AhGβ4*, and *AhGβ5* lines (Figure 2A-B). Similar partial complementation was observed in other developmental traits including leaf shape/area, silique length and dark-grown hypocotyl length. Overall, complete restoration, similar to WT, was not observed for any of the developmental traits (Supplementary Figure 5A-F).

**Figure 2.**
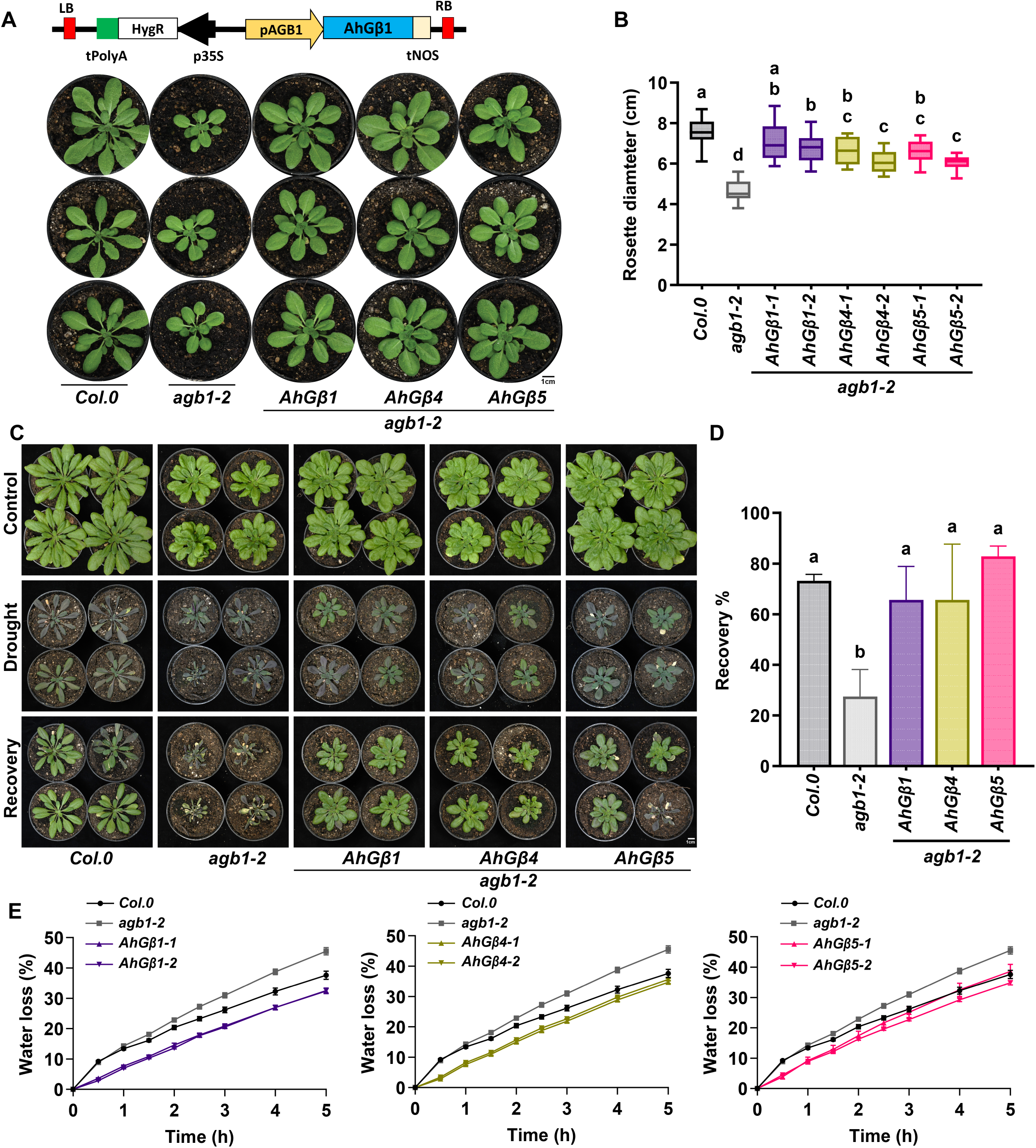
AhGβs complementation in Arabidopsis Gβ mutant *agb1-2* restored drought-sensitive phenotypes. (A) The diagram represents the T-DNA region of the complementation construct and the rosette morphology of *AhGβ* complementation lines with *agb1-2* mutant and wild-type (*Col.0*), scale 1 cm. (B) The rosette diameter measurement as a box plot. The middle line in the box plot represents the median, and the box range indicates the 25th to 75th percentiles. The whiskers denote the minimum-to-maximum range. The letters on top of boxes indicate the statistically significant differences (p<0.05, one-way ordinary ANOVA with Fisher’s LSD test). (C) The vegetative drought and recovery stages of *AhGβ* complementation lines with control plants, scale 1 cm. (D) The bar graph shows the percentage of recovery after the drought stress. The letters on top of boxes indicate the statistically significant differences (p<0.05, one-way ordinary ANOVA with Fisher’s LSD test). (E) The line graphs show the percentage water loss from detached rosettes of *AhGβ1, AhGβ4,* and *AhGβ5* complementation lines over time, compared with *agb1-2* and *Col.0.*

The *agb1-2* mutant is highly sensitive to vegetative drought stress due to higher water loss (Liu *et al*., 2017), making it an ideal system to evaluate whether *AhGβ* subunits can confer functional restoration under water-deficit conditions. The drought stress treatment on complementation lines resulted in complete rescue of drought sensitivity in *agb1-2* (Figure 2C). Compared to lowest recovery rate of *agb1-2* (∼27.5%), *AhGβ* complementation lines displayed significantly enhanced recovery rates ranging from ∼65% to 82%, which was more or less similar to WT recovery rate (∼73%), highlighting their functional conservation in drought stress adaptation (Figure 2D). As the drought sensitivity of the *agb1-2* is associated with higher water loss, we performed a longitudinal water-loss assay using detached rosettes to assess transpiration dynamics. The *agb1-2* mutant exhibited rapid water loss, losing ∼38% within 4 hours of detachment compared to ∼32% in WT. Interestingly, all *AhGβ* complementation lines exhibited significantly reduced water loss, with values ranging from ∼26% to 32% (Figure 2E). Previous studies have reported that *agb1-2* mutants are hypersensitive to ABA-induced inhibition of germination. In the presence of ABA, WT had ∼52.7% germination on the second day compared to *agb1-2* mutant which had a significantly lower germination rate (∼30.8%), whereas, all *AhGβ* complementation lines exhibited higher germination frequencies, ranging between 50% and 80% (Supplementary Figure 5G). Together, these findings indicate that *AhGβ* subunits not only partially complement the morphological defects of *agb1-2* but also entirely restore drought-tolerance by regulating water balance and stress physiology. Additionally, these results also suggested that all three AhGβ subunits are analogous when complemented in model plant Arabidopsis.

### 2.3. *AhGβ1* subunit positively influences plant growth and yield in groundnut

Both *AhGβ1* and *AhGβ3* were early drought-inducible genes, and *AhGβ1* functionally complemented drought-sensitive phenotype of *agb1-2*. Therefore, we generated stable transgenic overexpression (35Spro:*AhGβ1*) and an RNA interference (*RNAi*) knock-down line targeting primarily *AhGβ1* and *AhGβ3* transcripts (Figure 3A), to investigate the functional role of early stress-responsive Gβ subunits in groundnut. Putative transformants were confirmed using genotyping PCR using *NptII* and *BlpR* gene primers for overexpression and knock-down respectively (Supplementary Figure 6A-B). The relative expression of *AhGβ1* gene was significantly high in the overexpression lines (Figure 3B). In the knock-down lines, a 60-80% reduction in *AhGβ1* expression was observed, whereas only 18-50% reduction was observed for *AhGβ3* transcripts (Figure 3C, Supplementary Figure 6C). In addition, other *AhGβ* genes did not display a reduction in their expression, which was expected by the predicted number of efficient siRNAs (Supplementary Figure 6D). To elucidate the functional contribution of *AhGβ1* in plant development and yield, the transgenic lines were phenotypically evaluated for the following traits: plant height, total number of branches, peg number, pod number, seed number, seed size (width), seed weight and stomatal number. The overexpression lines consistently produced taller plants, with a mean height of ∼45 cm at 60 days after germination (DAG), compared to ∼40 cm in WT. In contrast, the *RNAi* lines showed a clear reduction, averaging at ∼37 cm (Figure 3D-E). A similar trend was observed for total branch number (including both primary and secondary branches), where *AhGβ1:Ox* displayed a higher branch number with a median of 9, compared to WT (median of 8). In contrast, *RNAi* lines exhibited a pronounced reduction, with a median branch number of 6 (Figure 3F). The number of pegs and pods serves as a critical determinant of reproductive success, as each peg represents a potential site for pod and seed development. The *RNAi* lines had the lowest peg number, with a mean of 64/plant, compared to 76/plant in WT; whereas the *AhGβ1:Ox* exhibited a prominent increase in peg number with a mean of 90/plant (Figure 3G). Similar pattern was observed in case of pod number, where *AhGβ1:Ox* possessed ∼40 pods/plant compared to WT (31/plant), and *RNAi* plants contained the lowest number of pods with a mean of 24/plant (Figure 3H). The number of pods directly correlated with the total number of seeds/plant, where *AhGβ1:Ox* displayed the highest number of seeds (fully developed) with an average of 48/plant. But *RNAi* exhibited the lowest number with an average of 24 seeds/plant compared to WT (32 seeds/plant) (Figure 3I). Previous reports in several crop species have highlighted the role of G-protein β subunits in regulating seed size and overall seed yield (Fan *et al*., 2006, Huang *et al*., 2009, Mao *et al*., 2010, Li *et al*., 2012, Vavilova *et al*., 2017, Sun *et al*., 2018b). Consistent with this, *AhGβ1:Ox* exhibited larger seed size/width (≈9 mm) compared to WT (≈7.7 mm), whereas *RNAi* lines produced overall smaller seed size with an average width of ≈6.8 mm (Figure 3J-K). The reduction in seed size also affected the 10 seed weight in *RNAi* lines, where an average of ≈2.6g was observed compared to ≈3.6g in WT and ≈4.15g in *AhGβ1:Ox* (Supplementary Figure 6E). Taken together, these results indicated that *AhGβ1* can contribute directly to vegetative growth, reproductive development and yield determination in groundnut, enhancing both peg initiation and successful seed setting when overexpressed, while their suppression can compromise these traits.

**Figure 3.**
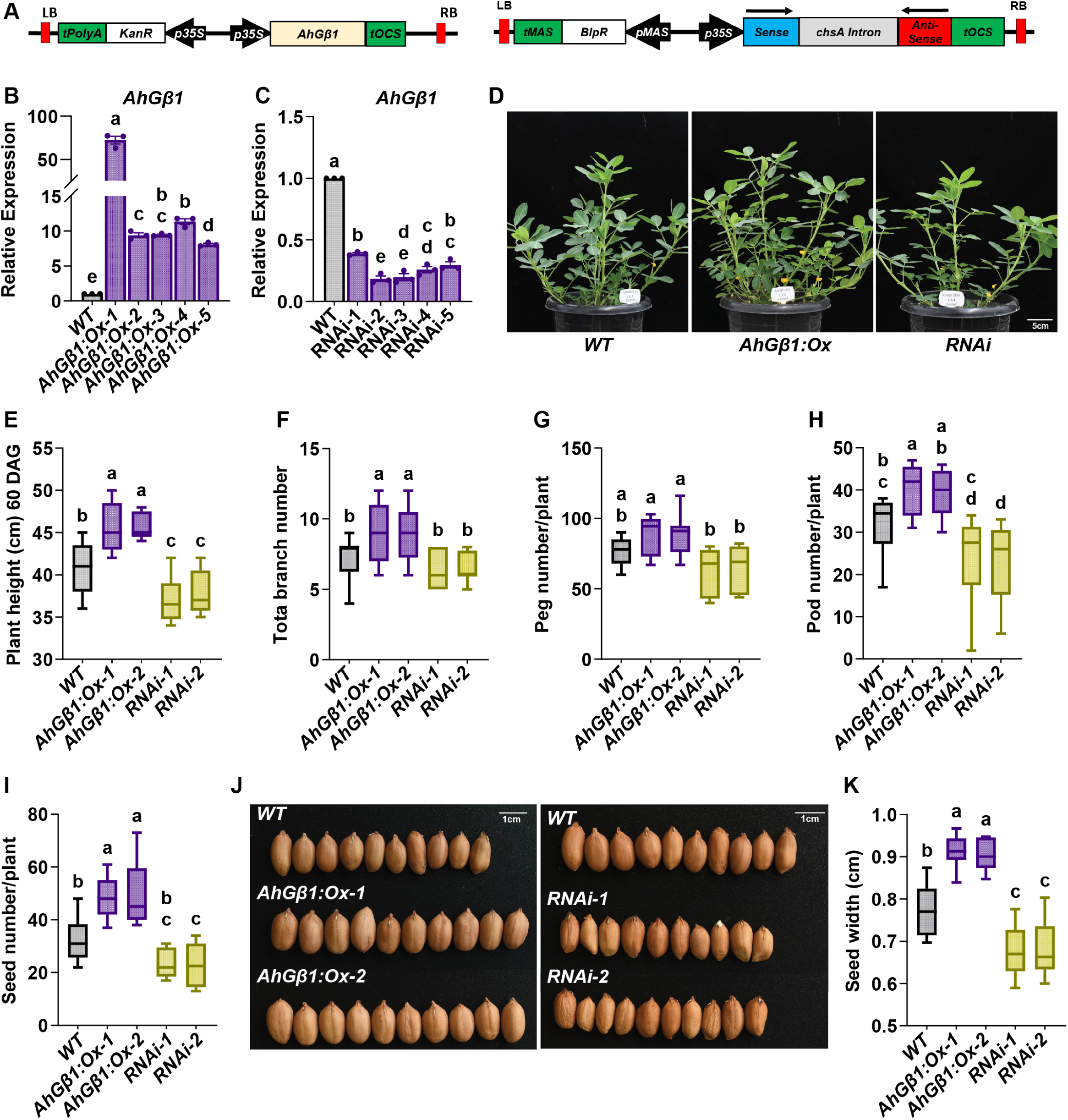
AhGβ1 overexpression enhances growth and yield in groundnut. (A) The illustration represents the T-DNA regions of overexpression and RNAi knockdown constructs. (B-C) The bar graphs represent the relative expression of the *AhGβ1* gene in the overexpression lines (B) and *RNAi* knockdown lines (C) in comparison to the wild-type (WT) control. The letters on top of boxes indicate the statistically significant differences (p<0.05, one-way ordinary ANOVA with Fisher’s LSD test). (D) The morphological differences *of AhGβ1* over-expression plants with *RNAi* knockdown and wild-type plants, photographs of 60 days old plants, scale 5 cm. (E-l) Box plots shows phenotypes across *WT, AhGβ1:Ox* and *RNAi* lines in control conditions, plant height (E), total branch number (F), peg number per plant (G), pod number per plant (H) and seed number per plant (I). (J) The seed-size morphology of the overexpression and knockdown lines compared to the wild-type control, scale: 1cm. (K) The box plot shows the seed width across *WT, AhGβ1:Ox* and *RNAi* lines in control conditions. The middle line in the box plots represents the median, and the box range indicates the 25th to 75th percentiles. The whiskers denote the minimum-to-maximum range. The letters on top of boxes/bars indicate the statistically significant differences (p<0.05, one-way ordinary ANOVA with Fisher’s LSD test).

### 2.4. *AhGβ1* subunit positively regulates drought-tolerance in groundnut

To determine the functional role of *AhGβ1* in mediating drought tolerance in groundnut, we performed a drought stress experiment using the transgenic lines followed by rehydration to assess recovery. During the stress period, *AhGβ1:Ox* plant displayed reduced wilting relative to WT, while *RNAi* lines exhibited severe wilting (Figure 4A). After rehydration, *AhGβ1:Ox* lines displayed a higher recovery rate of up to 95% compared to 70% recovery of WT, while *RNAi* lines displayed a lower (46%) recovery rate (Figure 4B). The phenotypic traits were evaluated during drought stress and in recovered plants. The *AhGβ1:Ox* lines displayed augmented height (∼38 cm) compared to WT (∼35 cm), while the *RNAi* lines displayed a significant reduction (∼30 cm) (Figure 4C). Branch number per plant was higher in *AhGβ1:Ox* lines (∼8-10 branches) compared to WT and *RNAi* lines (∼6-7) (Figure 4D). The peg number per plant after recovery was moderately increased in *AhGβ1:Ox* lines (∼75) compared to WT (∼60), whereas in *RNAi* lines a slight decrease was observed (∼50) (Figure 4E). The pods produced after the drought recovery were high in the *AhGβ1:Ox* lines (∼35 pods/plant) compared to WT (∼26 pods/plant), while the *RNAi* lines exhibited a pronounced reduction (∼16 pods/plant) (Figure 4F). Similar pattern was observed for the seeds produced per plant, where *RNAi* lines produced the fewest seeds (∼10/plant) in comparison to the WT (∼22/plant), while *AhGβ1:Ox* lines exhibited a significant increase in seed number after recovery (∼33/plant) (Figure 4G). Seed quality was similar to non-stress conditions observations, *AhGβ1:Ox* lines produced significantly larger seeds (∼8.8 mm width) compared with WT (∼7.2 mm), while *RNAi* lines showed reduced seed width (∼6.8 mm) (Supplementary Figure 7A). In terms of biomass, the 10-seed weight of *AhGβ1:Ox* lines averaged ∼3.5g, higher than WT (∼2.8g), whereas *RNAi* plants recorded the lowest (∼2.1g) (Supplementary Figure 7B). These findings collectively indicate that *AhGβ1* plays a central role in enhancing drought-tolerance in groundnut, not only by improving stress survival and recovery but also by sustaining developmental and yield-related traits under water-limiting conditions.

**Figure 4.**
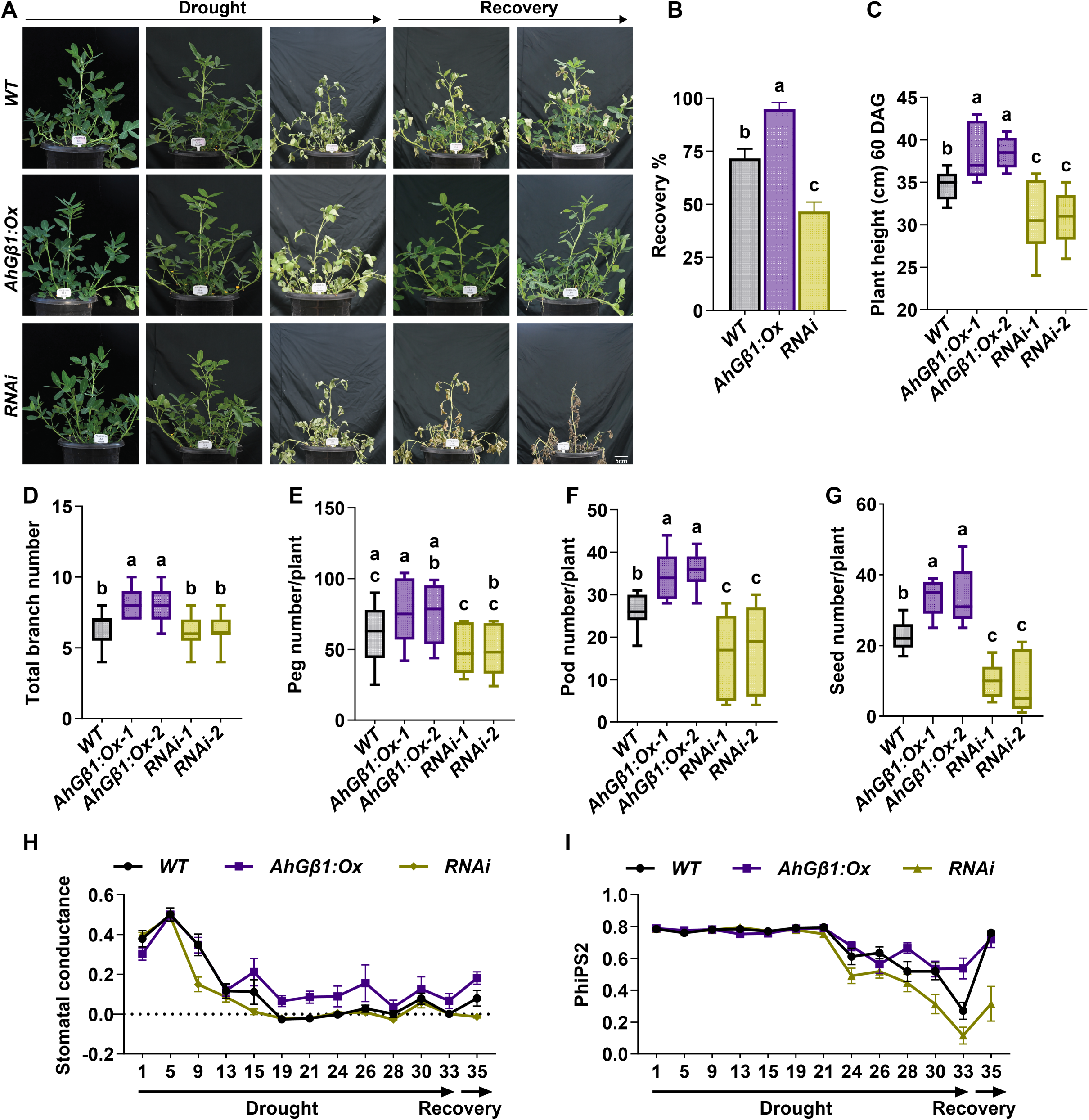
AhGβ1 overexpression improved drought-tolerance in groundnut. (A) The drought stress and recovery images of *AhGβ1* overexpression and knock-down lines with *wild-type (WT)* control, scale 5 cm. (B) The bar graph represents the percentage of recovery observed in the *AhGβ1* overexpression and *RNAi* knock-down lines with wild-type control. (C-D) Box plots represent phenotypes across *WT, AhGβ1 :Ox,* and *RNAi* lines under drought conditions for plant height (C) and total branch number (D). (E-G) Box plots represent phenotypes across *WT, AhGβ1 :Ox,* and *RNAi* lines after harvest (post-drought recovery): peg number per plant (E), pod number per plant (F), and seed number per plant (G). The middle line in the box plots represents the median, and the box range indicates the 25th to 75th percentiles. The whiskers denote the minimum-to-maximum range. The letters on top of boxes/bars indicate the statistically significant differences (p<0.05, one-way ordinary AN OVA with Fisher’s LSD test). (H-l) The line graphs show the stomatai conductance (H) and PhiPS2 (I) of *WT, AhGβ1:Ox,* and *RNAi* lines across the drought and recovery phases (X-axis denotes the days).

The physiological parameters monitored during the drought-stress period included stomatal conductance (g_sw_), total transpiration, and photochemical efficiency of photosystem II under light-adapted conditions (PhiPS2). Stomatal conductance (g_sw_) reflects the extent of gas exchange and indicates the degree of stomatal opening or closure. Under drought stress, plants tend to close their stomata and reduce water loss, enhancing survival. The stomatal conductance of all the lines showed a decline from 9^th^ to 13^th^ day of drought. Interestingly, the *AhGβ1:Ox* lines maintained a basal level of stomatal conductance (∼0.1-0.2) even during later stages of drought, whereas the stomatal conductance of WT and *RNAi* lines reached zero by 19^th^ day (Figure 4H). A similar pattern was observed for the total transpiration as well (Supplementary Figure 7C). The maintenance of basal stomatal conductance in *AhGβ1:Ox* lines was further supported by the PhiPS2 values, which remained in the intermediate range (∼0.3-0.5) during later stages of drought, while the *RNAi* lines displayed a huge decline compared to WT plants, substantiating their lower recovery (Figure 4I). Additionally, it has also been reported in Arabidopsis that *AGB1* can regulate stomatal development (Cao *et al*., 2021). Similar results were observed here as well, that *AhGβ1:Ox* displayed a reduction in stomatal number per unit area compared to higher stomatal number in WT and *RNAi* lines (Supplementary Figure 7D-E), which might contribute to reduced water loss in *AhGβ1:Ox* lines. These observations collectively support the higher growth and yield of the *AhGβ1:Ox* lines under drought stress conditions and signify their role in maintaining growth under stress.

### 2.5. Transcriptome analysis uncovers the role of *AhGβ1* in flavonoid biosynthesis and cell wall regulation under drought in groundnut

To understand the molecular mechanisms of *AhGβ1*-mediated drought-tolerance in groundnut, we have performed transcriptomic analysis using the *AhGβ1:Ox* and wild-type (WT) plants under drought and control conditions. Principal component analysis (PCA) revealed a non-overlapping cluster of samples, suggesting that conditions and genotypes significantly contribute to overall transcriptomic variability (Figure 5A). Differentially expressed genes (DEGs) were identified from three comparisons: *AhGβ1:Ox* versus WT in control (3446), *AhGβ1:Ox* versus WT in drought (2154) and drought versus control for WT (3133) (Figure 5B). By combining all three comparisons, a total of 7024 significantly altered transcripts were obtained (Supplementary Table 4), and subsequently used for k-mean clustering based on the z-score expression values. This resulted in eight different gene clusters with different expression patterns across the samples, and the functional analysis of this set of genes was displayed using KEGG and GO enriched terms (Supplementary Figure 9). Cluster 1 genes displayed a trend of a higher mean expression in WT control condition, while in WT drought, as well as in both *AhGβ1:Ox* conditions, the mean expression was low. Gene functional analysis revealed that these genes are majorly associated with cell wall-related terms and pathways, such as lignin catabolism, cell wall organization, apoplast, plant-type cell wall, etc. (Figure 5C). The qualitative difference in lignin and cellulose depositions further supported that *AhGβ1:Ox* lines have reduced cell wall component accumulation compared to WT and *RNAi* lines (Supplementary Figure 10A-B). The Calcofluor White (CFW) fluorescence quantification displayed a significant reduction in *AhGβ1:Ox* line in comparison to WT, whereas the *RNAi* lines showed similarity to WT level CFW fluorescence (Supplementary Figure 10C). Further, downregulation of cellulose synthase A (CESA), CESA-like (CSL), beta-xylosidase (XYL4), etc. gene families in *AhGβ1:Ox* lines proposed the possible involvement of *AhGβ1* in cell wall biogenesis regulation (Supplementary Figure 11A-B; 12A-C).

**Figure 5.**
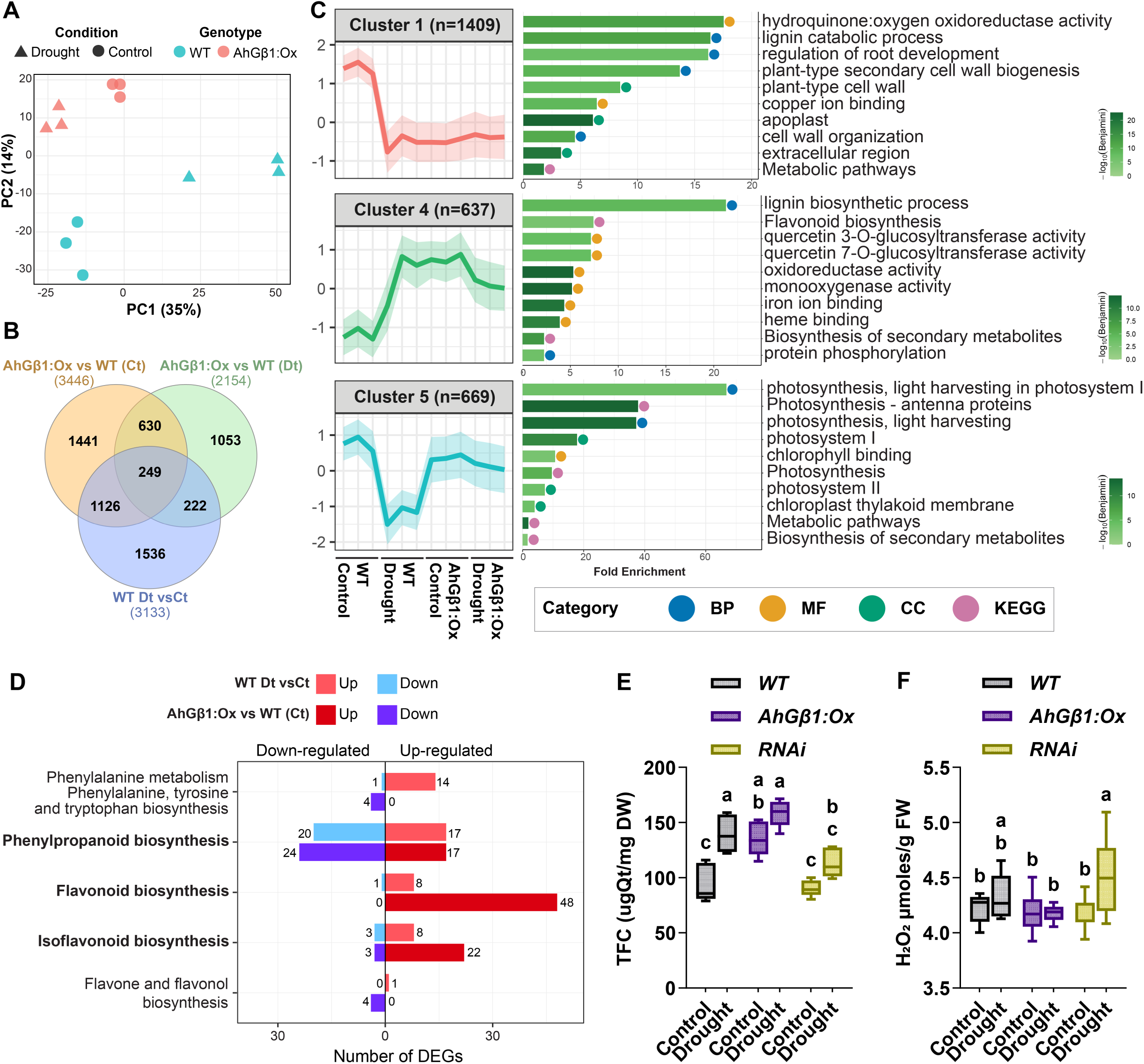
Transcriptional regulation of cell wall organization and flavonoid biosynthesis by *AhGβ1* under drought stress. (A) The PCA plot shows the overall variation in expression across genotypes and conditions. (B) The Venn diagram shows the overlapping DEGs across the comparisons, and the bracketed numbers represent the counts of DEGs in each comparison. (C) The line graphs show the average expression curves for k-mean clusters across samples, along with the number of genes in each cluster. The bar graphs are showing the top 10 GO terms or KEGG pathways of the corresponding cluster, bar colour represents the significance as -log10(Benjamini). The bubble colour represents the category of term/pathway (BP-biological process, MF-molecular function, CC-cellular component and KEGG pathway). All eight clusters are represented in Supplementary Figure 9. (D) The mirror bar graphs represent the number of DEGs in the corresponding biosynthetic pathways for two different comparisons. Color represents up- and down-regulated genes of the corresponding comparison. (E-F) The box plots show the total flavonoid content (E) and H2O2 levels (F) in the *WT, AhGβ1:Ox* and *RNAi* lines under control and drought conditions. The middle line in the box plots represents the median, and the box range indicates the 25th to 75th percentiles. The whiskers denote the minimum-to-maximum range. The letters on top of boxes/bars indicate the statistically significant differences (p<0.05, one-way ordinary ANOVA with Fisher’s LSD test).

An interesting pattern observed in cluster 4, was the increase in the mean transcript abundance of genes in drought condition of WT vs control, while a similarity in transcript abundance of *AhGβ1:Ox* in control to that of WT drought and moderately reduced in *AhGβ1:Ox* in drought condition. These genes were primarily mapped to crucial secondary metabolism like flavonoid and lignin biosynthesis, quercetin 3-O-glucosyltransferase activity etc., which are related to formation of antioxidant compounds, a prerequisite for abiotic stress tolerance (Figure 5C). This pattern demonstrated that these genes might be vital for drought-tolerance in WT, and AhGβ1 might be imperative for regulating their expression.

Similar enrichment of genes involved in flavonoid and other secondary metabolites biosynthesis was observed in clusters 3 and 6 (Supplementary Figure 9). These two clusters showed the maximum mean expression in the control condition of *AhGβ1:Ox*, signifying the possible role of *AhGβ1* in the regulation of secondary metabolism. Whereas, the expression declined in *AhGβ1:Ox* drought suggesting a possible negative feedback/homeostatic regulation by the formation of intermediary compounds in these pathways. Further we analyzed the number of DEGs associated with phenylpropanoid, flavonoid, and isoflavonoid biosynthesis (Figure 5D). In WT drought plants compared to the control condition, 37 DEGs were observed in phenylpropanoid biosynthesis, including 17 upregulated and 20 downregulated (Figure 5D). This pathway can contribute to both lignin and flavonoid biosynthesis. In *AhGβ1:Ox* plants compared to WT in control conditions, 41 DEGs were mapped to the same pathway, with 17 upregulated and 24 downregulated genes. These included key enzymes such as: phenylalanine ammonia-lyase (PAL), cinnamyl-alcohol dehydrogenase (CAD), cinnamoyl-CoA reductase (CCR), peroxidase (PER), etc., further supporting that this pathway is differentially regulated under drought in WT and *AhGβ1:Ox* lines (Supplementary Figure 11C; 12D-F). In the flavonoid biosynthesis pathway, 8 DEGs were upregulated in WT plants under drought stress relative to controls. In contrast, *AhGβ1:Ox* plants showed a much stronger response, with 48 upregulated DEGs compared with WT controls. These included key flavonoid biosynthetic genes such as chalcone synthase (CHS), chalcone isomerase (CHI), and chalcone reductase (CHR) (Figure 5D; Supplementary Figure 11D; 12G-I). Similarly, isoflavonoid biosynthesis pathway showed higher activation in *AhGβ1:Ox* plants, where 22 DEGs were upregulated relative to WT controls. In WT drought plants relative to control, only 8 genes in this pathway were upregulated (Figure 5D, Supplementary Figure 11D; 12J). Gene regulatory network analysis predicted that *AhGβ1*-responsive transcription factors, including members of the MYB, bHLH, WRKY, ERF and NAC families, are associated with the regulation of phenylpropanoid and flavonoid biosynthetic genes (Supplementary Figure 13A-B). Together these results demonstrated that *AhGβ1* is imperative for the transcriptional regulation of biosynthetic pathways that lead to the production of antioxidant secondary metabolites.

The transcripts in cluster 5 showed higher expression in control condition for both WT and *AhGβ1:Ox*, while upon drought in WT the transcripts displayed a reduced expression, but not in *AhGβ1:Ox*, where the expression remains same as control. Most of these genes were associated with photosynthesis-related terms such as photosynthesis - antenna proteins, photosynthesis, chlorophyll binding, photosystem I, chloroplastic thylakoid membrane, etc. (Figure 5C). These expression profile indicated that *AhGβ1:Ox* signaling facilitates to protect/maintain photosynthetic machineries during drought stress compared to WT (Figure 5C). Collectively, these results indicated that drought stress suppresses cell wall-related process, and promotes the accumulation of anti-oxidant secondary metabolites such as flavonoids by regulating phenylpropanoid biosynthesis and other flavonoid biosynthetic pathways. This, in turn, protects the photosynthetic machineries, and *AhGβ1* might have a crucial role in the regulation of these pathways.

Further to validate our hypothesis, we analyzed the total flavonoid content (TFC) in the transgenic lines under both control and drought conditions (from 9^th^ day of drought) (Figure 5E). Upon drought stress in WT, the mean TFC level significantly increased from 93 μg Qt/mg to 139 μg Qt/mg. Interestingly *AhGβ1:Ox* lines displayed almost similar levels of TFC, like WT drought, under control conditions (134 μg Qt/mg), and upon drought, a less significant increase was observed (158 μg Qt/mg). The *RNAi* lines showed 90 μg Qt/mg mean TFC under control conditions, whereas upon drought, the increase was less notable (112 μg Qt/mg) (Figure 5E). A less significant increase of H_2_O_2_ levels was noticed in WT upon drought, whereas in the *AhGβ1:Ox* lines, no change was observed upon drought. As expected, *RNAi* lines displayed a significant increase in H_2_O_2_ levels upon drought (Figure 5F). Taken together, these results further confirmed that *AhGβ1*-mediated flavonoid biosynthesis contributes to the antioxidant system that mitigates the ROS produced during drought stress, thereby improving tolerance.

### 2.6. Metabolomic analysis confirms the role of *AhGβ1* in flavonoid biosynthesis under drought in groundnut

To corroborate the AhGβ1-mediated secondary metabolic regulations identified in transcriptome studies, we performed metabolomics analysis using drought and control samples from *AhGβ1:Ox* line and WT plants. The PCA plot highlighted pronounced clustering and clear differentiation between genotypes and conditions (Figure 6A). The *AhGβ1:Ox* control and drought samples displayed comparatively lesser variations, suggesting the overexpression-mediated accumulation of metabolites irrespective of condition (Figure 6A). The identified compounds were mainly distributed across different secondary metabolite sets such as phenylpropanoids (17), flavonoids (16), alkaloids (8), terpenoids (8), etc. and displayed a higher accumulation in the *AhGβ1:Ox* lines (Figure 6B). The differentially abundant metabolite (DAM) analysis was performed from three comparisons; *AhGβ1:Ox* versus WT in control (58), *AhGβ1:Ox* versus WT in drought (38) and drought versus control for WT (49) (Supplementary Figure 14A-B). Most of the phenylpropanoids and flavonoids were significantly abundant in the *AhGβ1:Ox* lines compared to WT in control conditions, and these compounds were abundant in WT drought conditions compared to control (Supplementary Figure 14B). Further, pathway enrichment analysis also confirmed significant regulation of the flavanone and flavonol, phenylpropanoid, phenylalanine, tyrosine and tryptophan biosynthesis, etc. pathways during drought in WT and under control condition in *AhGβ1* overexpression lines (Figure 6C). The compounds detected in the above secondary metabolites biosynthesis pathways were plotted with their respective KEGG pathways. Most of these compounds displayed a higher abundance in control condition of *AhGβ1:Ox*, followed by *AhGβ1:Ox* drought and WT drought conditions. The metabolite output of these pathways were potent antioxidant compounds (Figure 6D, Supplementary Figure 14-18) like quercetin, rutin, baimaside, etc. which were highly abundant in the *AhGβ1:Ox* plants, whereas during drought, a slight reduction was observed for some metabolites (Supplementary Figure 14C-D). Collectively, these results further confirmed that *AhGβ1* acts as an upstream regulator of these pathways to produce the antioxidant flavonoid compounds to mitigate the drought stress. The overexpression of *AhGβ1* leads to upregulation of genes and accumulation of these compounds, and that the reduction in expression of these pathway enzymes and metabolite abundance during drought in *AhGβ1:Ox* lines might be connected to the negative feedback/homoeostasis regulation (Supplementary Figure 15-18).

**Figure 6.**
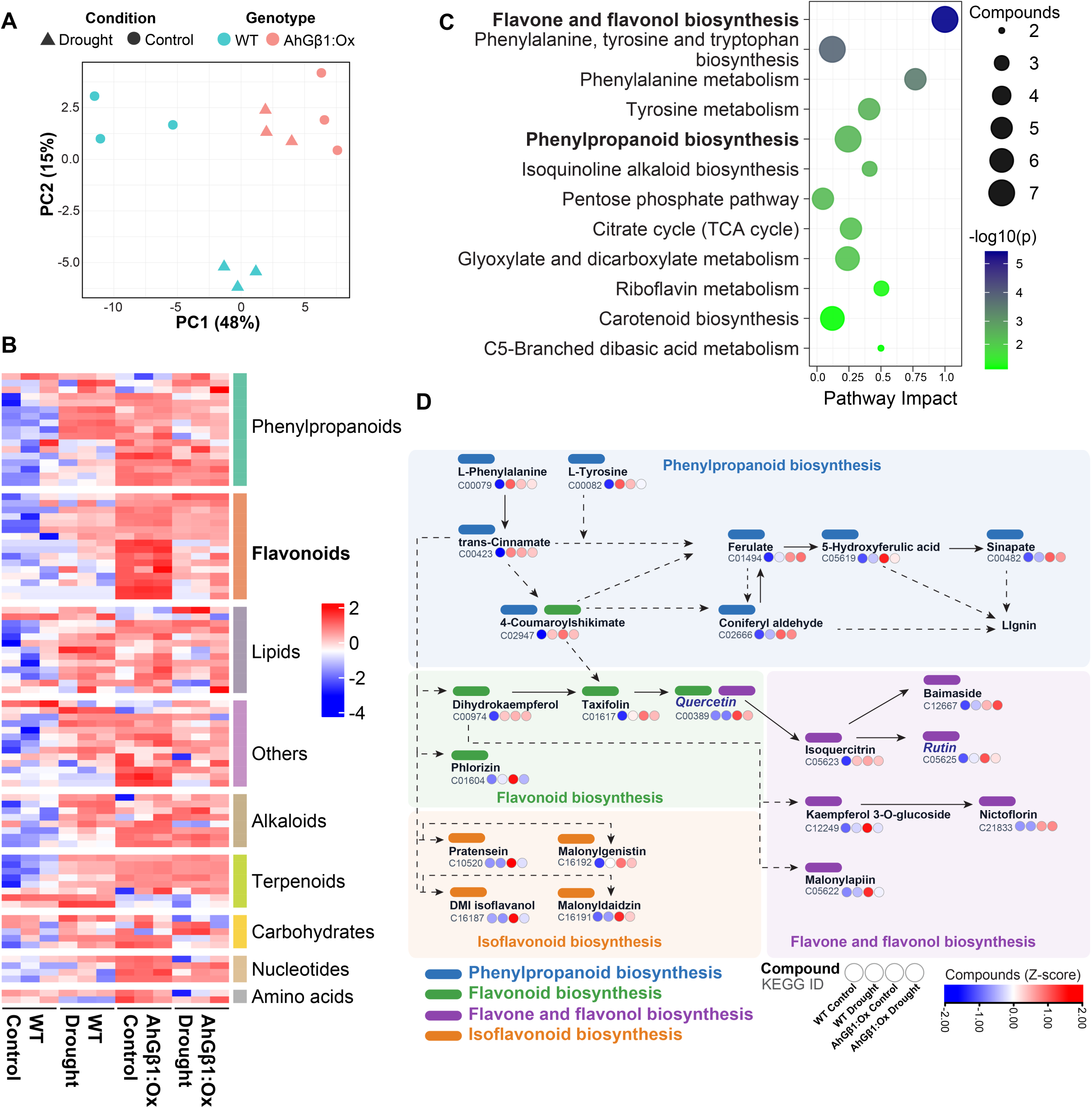
*AhGβ1* overexpression promotes the accumulation of drought-induced flavonoid metabolites. (A) The PCA plot exhibits the variation in metabolite intensities across genotypes and conditions. (B) The heatmap shows the Iog2-transformed raw intensity values for different metabolite classes across control and drought samples from *WT* and *AhGβ1:Ox* lines. The colour scale represents the Iog2-transformed raw intensity values. (C) The bubble plot represents the KEGG pathway enrichment analysis, X-axis represents the pathway impact, bubble size denotes the number of compounds, and bubble colour denotes the significance (-log10(pvalue)). (D) The pathway diagram represents the compounds identified across the phenylpropanoid, flavonoid, Isoflavonoid, flavone and flavonol biosynthesis pathways. The colour bubbles represent the mean intensity values of compounds across the samples, colour scale denotes the Z-score abundance. The colour bar on top of the compound denotes the KEGG pathway. Line arrows represent the direct reaction/ pathway connection, and dashed line arrows denote indirect pathway connections.

### 2.7. Treatment with *AhGβ1*-regulated flavonoids quercetin and rutin, enhances drought-tolerance in groundnut

To determine the potential antioxidant activity of AhGβ1-regulated flavonoid compounds, we selected bioactive quercetin and its storage form, rutin, as the potential candidates, and analyzed drought recovery, physiological parameters, and H_2_O_2_ content following their foliar application onto WT plants. Quercetin hydrate (150 μM) and rutin hydrate (0.5 mM) were sprayed on alternative days as treatments and DMSO (0.1%) was sprayed as control, followed by 27 days of drought stress (D) and five days of recovery (R). The quercetin and rutin-treated plants displayed reduced wilting symptoms compared to DMSO-treated plants under drought, and the corresponding plants under control condition exhibited better growth (Supplementary Figure 19A). The recovery percentage further showed that quercetin and rutin treatments enhanced drought-tolerance in groundnut with 70% and 55% recovery, respectively, compared to 0% in DMSO treatment (Figure 7A). The physiological parameters such as stomatal conductance (g_sw_), total transpiration, and photochemical efficiency of Photosystem II under light-adapted conditions (PhiPS2) were enhanced in the control condition of the quercetin and rutin-treated plants compared to DMSO-treatment. During drought stress, g_sw_ declined faster in DMSO-treated plants by day 10 and failed to recover, whereas the rutin-treated plants declined to lower limit by day 15 and recovered after rewatering (0.04-0.06). Interestingly the quercetin-treated plants maintained a lower level of conductance until day 25 (0.05-0.01) and recovered after rewatering (0.2-0.4) compared to DMSO-treated plants (0.0) (Figure 7B). A similar pattern was observed in the case of total transpiration as well (Supplementary Figure 19B). The PhiPS2 also declined faster in the DMSO-treated plants during drought, whereas quercetin and rutin-treated plants retained a reduced level (0.1-0.3) and improved after recovery (0.4-0.6) (Figure 7B). These results indicated that these antioxidant compounds protect the photosynthetic machineries and sustain their function during drought. Confocal imaging of DPBA (a flavonol-binding dye)-stained epidermal peels revealed fluorescence signals predominantly in the stomatal guard cells. Plants treated with quercetin and rutin displayed increased flavonol-associated fluorescence compared with the DMSO control (Figure 7C; Supplementary Figure 19C). Consistent with these observations, biochemical quantification of total flavonoid content in the leaves showed a similar pattern of accumulation (Figure 7D). Together, these results indicate that exogenous quercetin and rutin treatments increased flavonoid/flavonol-associated metabolite abundance in groundnut leaves. To further validate the amount of ROS produced during drought, we measured H_2_O_2_ levels. The DMSO-treated plants displayed a huge increase under drought (8.5 µmoles/g increase), whereas the quercetin and rutin treated plants displayed a marginal reduction in the control condition itself and a minimal increase was observed upon drought compared to DMSO-treated plants (2.4-3.2 µmoles/g increase) (Figure 7E). These results further confirmed that the drought-induced ROS potentially damages the guard cells and photosynthetic apparatus in DMSO-treated plants, leading to their death. On the other hand, antioxidant compound treatment effectively suppresses the formation of the ROS, protects membranes and chloroplasts, and maintains little uptake of CO_2_ and photosynthesis even under the drought stress condition, subsequently contributing to better tolerance and recovery. Overall, the application of such antioxidant compounds enhances the growth, yield and drought-tolerance in groundnut, and this can be considered as a prospective application in large-scale production in fields as well.

**Figure 7.**
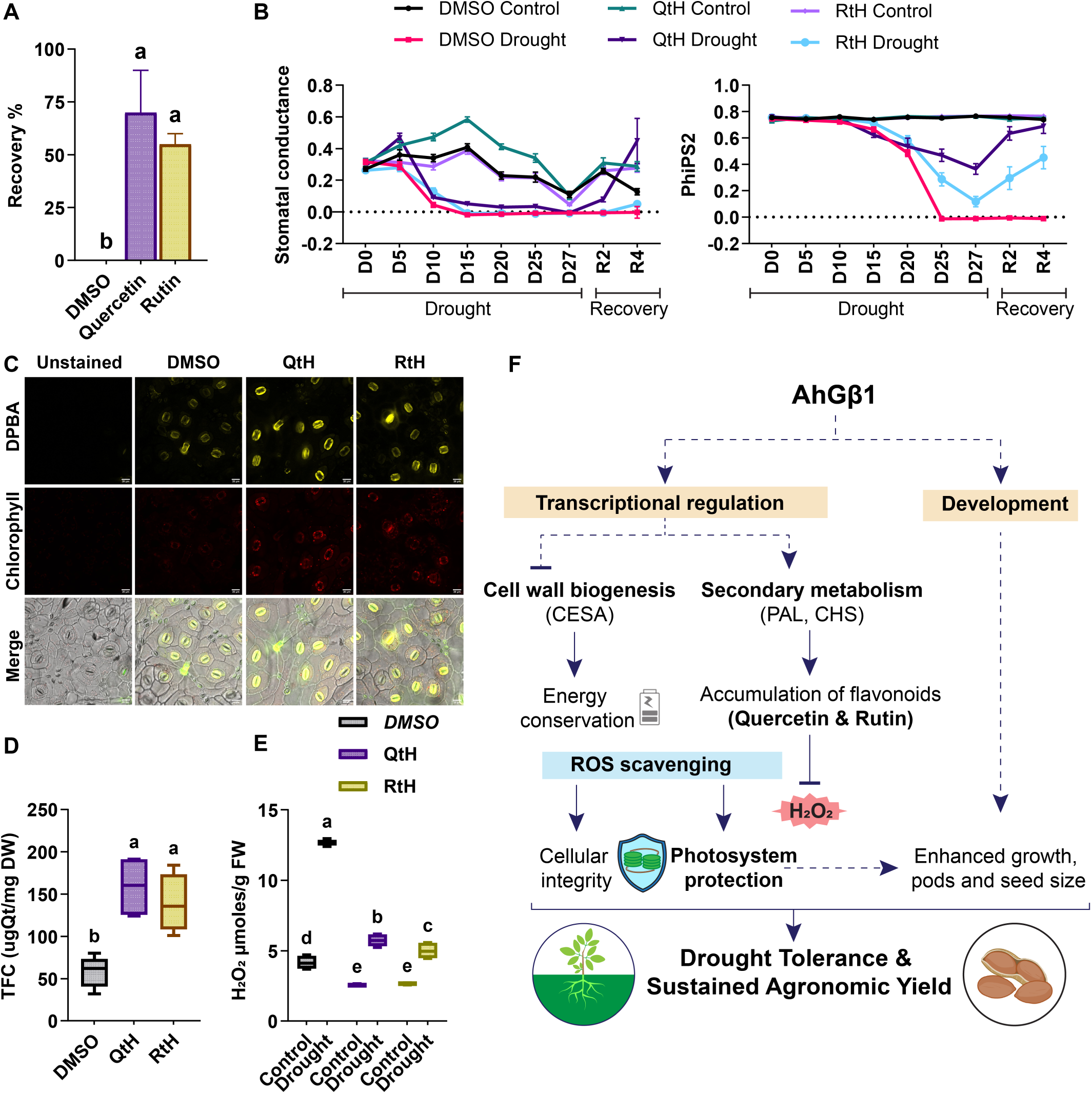
Quercetin and rutin treatments improved drought-tolerance in groundnut. (A) The bar graph shows the percentage recovery observed in plants treated with DMSO, quercetin, and rutin after drought. (B) Line graphs display the stomatai conductance and PhiPS2 across compound treated plants during drought stress (D0-D27), and recovery (R2 and R4) days. Line colour denotes the control and drought plants of DMSO, quercetin (QtH), and rutin (RtH) treatments. (C) Representative confocal images of DPBA-stained groundnut leaf epidermis after 5 days of foliar treatments. DPBA-flavanol fluorescence was represented in yellow. Chlorophyll autofluorescence was represented in red. Both merged with bright-field represented as Merged, Scale bar 20 pm. (D) The box plots show the total flavonoid content after the DMSO, quercetin and rutin treatments. (E) The box plot represents the H_2_O_2_ content in the plants under control and drought conditions of DMSO, quercetin and rutin treatments. The middle line in the box plots represents the median, and the box range indicates the 25th to 75th percentiles. The whiskers denote the minimum-to-maximum range. The letters on top of boxes indicate the statistically significant differences (p<0.05, one-way ordinary ANOVA with Fisher’s LSD test). (F) Schematic model illustrating the proposed mechanism by which *AhGβ1* enhances drought tolerance. AhGβ1 transcriptionally regulates genes involved in cell wall biogenesis and secondary metabolism, thereby suppressing cell wall biogenesis and promoting biosynthesis of antioxidant flavonoids. Accumulation of quercetin and rutin enhances ROS scavenging and lowers H_2_O_2_ levels. This facilitates the protection of photosystems and maintains cellular integrity under drought stress. Simultaneously, AhGβ1 regulates developmental pathways to promote plant growth, pod development, and seed size. These integrated physiological and developmental responses collectively provide enhanced drought tolerance and sustained agronomic yield.

## 3. Discussion

Drought stress can cause up to 85% reduction in groundnut yield (Reddy *et al*., 2003). As a major oilseed crop and dryland legume, groundnut productivity under drought stress represents one of the major agronomic challenges in semi-arid regions, where water scarcity progressively threatens yield and quality. Drought-tolerance in plants is a complex trait governed by multiple, interlinked mechanisms involving signaling networks, transcriptional regulation, and metabolic adjustments (Takahashi *et al*., 2018; Takahashi *et al*., 2020). In plants, heterotrimeric G-proteins act as the molecular switches that transduce extracellular signals from receptor-like kinases to intracellular effectors, regulating nearly all aspects of plant growth, development and stress responses (Pandey, 2019; Urano *et al*., 2013). In the current study, through a meta-transcriptome analysis we found tandemly duplicated *AhGβ1* and *AhGβ3* genes as early signaling components that might contribute to groundnut drought-tolerance. This aligns with the evidences from various plant species where Gβ subunit act as the master regulator of abiotic stress signaling, hormonal cross talk and developmental plasticity (Wang and Botella, 2022; Zhang *et al*., 2021).

### AhGβ subunits contributes to development and drought stress tolerance in groundnut

The functional characterization demonstrated that AhGβs are essential for both vegetative growth and reproductive success. The overexpression enhanced height, branching, peg initiation, pod number, and seed size, whereas *RNAi* knock-down severely compromised these traits. A few similar developmental regulations of G-protein subunits have been reported in model and crop plants. The Arabidopsis *agb1* mutant displayed reduced organ size and altered leaf shape (Wang and Botella, 2022). The overexpression of *AGG3* (Arabidopsis G-gamma 3) in *Camelina sativa* increased seed size, yield and improved stress tolerance (Roy Choudhury *et al*., 2014). In tomato the Gβ subunit (SlGB1) CRISPR knock-out mutant displayed reduced fruit and seed size (Wang *et al*., 2025). Moreover, in rice, the RGB1 (Rice G-beta 1) positively regulated grain size and ABA biosynthesis, acting as a positive regulator of drought-tolerance (Sun *et al*., 2018; Zhang *et al*., 2015).

The *AhGβ1:Ox* groundnut plants exhibited enhanced drought resilience characterized by reduced wilting, minimized water loss and higher recovery. Heterologous expression of *AhGβ* subunits in the drought-sensitive Arabidopsis *agb1-2* mutant completely restored drought-tolerance and complemented the transpirational water loss. The Gβ subunit is essential for abscisic acid (ABA)-mediated stomatal closure and the regulation of ion channels to prevent excessive transpiration and preserve turgor pressure (Fan *et al*., 2008; Wang *et al*., 2001). The Gβ subunit also physically interacts with receptor-like kinases FERONIA (FER), to regulate stomatal movement and ABA responses (Yu *et al*., 2018). Consistent with this conserved role, *AhGβ1:Ox* lines maintained minimum stomatal conductance and sustained photosystem II efficiency (PhiPS2) during drought. The enhanced drought tolerance and growth in *AhGβ1:Ox* suggested that they appear to function at the intersection of developmental and stress-responsive pathways. This dual ability highlights that G-proteins can govern developmental plasticity to ensure reproductive success and fitness under water-deficit conditions.

### AhGβ1 regulates cell wall organization and flavonoid metabolism under drought stress

A major mechanism underlying AhGβ1-mediated drought-tolerance is the extensive transcriptional reprogramming of cellular metabolism (Supplementary Figure 13A-B). Transcriptome analysis of *AhGβ1:Ox* plants under drought stress revealed a noticeable downregulation of genes associated with cell wall organization and biogenesis, such as cellulose synthases (CESA) and pectate lyases (PYL). Under drought stress, cell wall expansion becomes an energetically wasteful process due to reduced turgor pressure (Alvarez *et al*., 2008; Bray, 2004; Iraki *et al*., 1989; Piro *et al*., 2003; Sweet *et al*., 1990). Therefore, suppressing this energy and carbon-intensive process of cell wall expansion is a known stress-avoidance strategy to conserve metabolic resources (Han *et al*., 2025; Martínez *et al*., 2007; Zhu *et al*., 2023). Recent functional studies in tomato Gβ (*SlGB1*) have linked G-proteins to cell wall-related metabolite levels (Wang *et al*., 2025).

Our study demonstrated that AhGβ1 acts as a molecular switch to upregulate secondary metabolic pathways, including the biosynthesis phenylpropanoids, flavonoids, and isoflavonoids. Overexpression lines showed upregulated expression of key biosynthetic enzymes, including phenylalanine ammonia-lyase (PAL) and chalcone synthase (CHS). The regulation of these specific secondary metabolic pathways by G-proteins has been documented in other species. The GPA1 and GCR1 are known to mediate flavonoid biosynthesis and regulate stress-responsive transcriptional networks (Chakraborty *et al*., 2015). In tomato, Gβ signaling regulates phenylpropanoid pathways and secondary metabolism via MYB and bHLH transcription factors (Wang *et al*., 2025). Cumulatively, this metabolic rerouting suggests that AhGβ1 coordinates a strategic shift from growth-oriented cell wall organization to defense-oriented secondary metabolite production.

Drought-induced stomatal closure restricts CO_2_ uptake and leads to reactive oxygen species (ROS) production, which causes oxidative damage in cell membranes and to the photosynthetic machinery (Chaves *et al*., 2009; Miller *et al*., 2010). To overcome these plants produce ROS-scavenging enzymes (e.g., SOD, CAT, and APX) alongside non-enzymatic antioxidants like flavonoids (Hasanuzzaman *et al*., 2020). Our metabolomic profiling identified specific antioxidant flavonoids quercetin and rutin, which were highly abundant in *AhGβ1:Ox* lines. Subsequently, *AhGβ1:Ox* plants maintained stable, low H_2_O_2_ levels under drought stress in contrast to the high ROS accumulation and oxidative damage observed in *RNAi* lines. G-proteins regulate the activity of plasma-membrane-localized NADPH oxidases (RbohD and RbohF) which are required for ROS production and ABA-mediated stomatal closure (Kwak *et al*., 2003; Torres *et al*., 2013). Furthermore, overexpression of *Gβ* subunits augments drought-tolerance by enhancing ROS detoxification in mulberry and wheat (Liu *et al*., 2017; Xiong *et al*., 2023). Further, exogenous application of AhGβ1-regulated flavonoids (quercetin and rutin) enhanced drought recovery in wild-type groundnut plants. These treatments lowered H_2_O_2_ accumulation by higher flavonoid content in guard cells and preserved higher photosynthetic efficiency, effectively mimicking the drought-tolerant phenotype of the *AhGβ1:Ox* lines.

In conclusion, this study establishes that Gβ subunits are conserved regulators of developmental and stress-adaptive processes in plants. In groundnut, AhGβ1 regulates stress response by dynamically shifting metabolic flux away from energy-consuming cell wall biosynthesis towards enhanced accumulation of antioxidant flavonoids. This metabolic reprogramming effectively neutralizes drought-induced oxidative stress, safeguarding the photosynthetic apparatus and preserving cellular integrity. The functional conservation of *AhGβ1* and its impact on yield under stress conditions underscore their value as a molecular target for crop improvement (Figure 7F). The identification of AhGβ1-regulated protective metabolites such as quercetin and rutin not only advances our mechanistic understanding of G-protein signaling but also offers a potential agrochemical strategy, such as targeted foliar applications, to directly mitigate the impact of drought and sustain yield in groundnut cultivation.

## 4. Materials and methods

### 4.1 Meta-analysis of groundnut drought stress datasets

Four drought-stress transcriptome datasets of groundnut were retrieved from the SRA database (Bioproject: PRJNA687542 (Wang *et al*., 2021), PRJNA657965 (Jiang *et al*., 2021), PRJNA706902 (Zhao *et al*., 2021) and PRJNA629665 (Huang *et al*., 2020)). After initial analysis (Method S1) meta-analysis was conducted using the metaRNASeq R-package (Rau et al., 2014), Fisher’s combined probability test was applied to combine pvalues across the comparisons. Genes were retained if they showed significant meta-analysis results (adjusted p-value<0.05), and exhibited |log2 fold-change|≥1.5 at least in 4 comparisons for early and 5 comparisons for late. Functional gene enrichment analysis was performed using DAVID Bioinformatics tool and Mercator4 (V8.0) annotation tools.

### 4.2 Stress treatment and expression studies of AhGβs

Two-week-old *Arachis hypogaea* seedlings were subjected to treatments with 100 µM abscisic acid (ABA), 20 % polyethylene glycol (PEG), and 200 mM mannitol. Leaf samples were collected at 1 h, 12 h, and 24 h post-treatment and immediately frozen in liquid nitrogen. Drought stress was imposed at flowering stage by withholding the water for 30 days. Leaf samples were collected different days (1, 3, 5, 13, 19 and 21) and frozen in liquid nitrogen. RT-qPCR was performed using Method S2.

### 4.3 Plant materials - Arabidopsis complementation lines

The complementation lines were generated in the *Arabidopsis thaliana agb1-2* mutant background (T-DNA insertion line; ABRC stock number CS6536). For the complementation constructs, a 1.5 kb fragment of native *AtAGB1* promoter was PCR-amplified from wild-type Col-0 genomic DNA using primers containing EcoRI and NcoI restriction sites (Supplementary Table 5) and cloned into pCAMBIA1305.2 vector. Full-length coding sequences of AhGβ1, AhGβ4, and AhGβ5 were amplified with primers having XbaI and BstEII sites and cloned downstream of the *AtAGB1* promoter within the same vector using restriction-ligation cloning. *Agrobacterium tumefaciens* strain GV3101 carrying the constructs was used to transform *agb1-2* plants via the floral dip method (Zhang *et al*., 2006), and homozygous T3 and T4 lines were confirmed and used for subsequent studies. Phenotyping and drought experiments were performed as per Method S5.

### 4.4 Generation of AhGβ over-expression and knock-down lines in groundnut

The full-length coding sequences of *AhGβ1* were PCR-amplified using gene-specific primers containing XbaI and BstEII restriction sites (Supplementary Table 5) and cloned into pCAMBIA2301 binary vector in which the 35S promoter had been inserted into the multiple cloning site to drive constitutive expression. For functional knock-down of *AhGβ1* and *AhGβ*3, a 300 bp conserved region derived from *AhGβ1* was selected as an RNA interference (*RNAi*) trigger sequence to target both *AhGβ* homologs. The target region was identified using SiFi21 software with default parameters to ensure maximum on-target efficiency and minimal off-target effects (Supplementary Figure 6D). The 300 bp fragment was PCR-amplified using a forward primer carrying XhoI-XbaI sites and a reverse primer with BamHI-NcoI sites (Supplementary Table 5). The amplified fragment was sequentially cloned into the pFGC5941 vector. The *Agrobacterium tumefaciens* strain C58C1 carrying the construct was used for plant transformation. Genetic transformation of *Arachis hypogaea* was performed using the tissue culture method optimized for cotyledonary explants (Sharma and Anjaiah, 2000). For screening the putative transgenic lines, genomic PCR was performed with gene-specific primers targeting the *NptII* selectable marker for overexpression lines and the *Bar* gene for *RNAi* knock-down lines. In addition, transcript-level changes were also confirmed in these lines using qRT-PCR with gene-specific primers for *AhGβ* genes. Phenotyping and drought stress experiment using transgenic groundnut lines was performed as per Method S6.

### 4.5 Transcriptome analysis

Leaf samples were collected on the fifth day of drought stress from control and drought treatment plants in triplicate for RNA sequencing. After initial analysis using Method S7, protein-coding gene types were further utilized for downstream analysis (Supplementary Figure 8). The k-mean clustering was performed in R. The GO and KEGG analyses were performed using DAVID Bioinformatics tool (https://davidbioinformatics.nih.gov/tools.jsp). An integrated transcription factor-target gene regulatory network with correlation was generated using the Method S8.

### 4.6 Untargeted metabolome analysis

Leaf samples from fifth day of drought stress with controls, were used for metabolite extraction. Untargeted metabolomic profiling was conducted using UHPLC coupled with high-resolution mass spectrometry (HRMS) on an Agilent 1290 Infinity II LC system interfaced with an Agilent 6545XT Q-TOF mass spectrometer. After initial analysis (Method S9) differentially abundant metabolite (DAM) analysis was performed using the limma R-package, and significant metabolites were defined using |log2FC|>1 and FDR<0.05. Metabolite identification was performed using exact mass and MS/MS fragmentation patterns through Metaboanalyst (Pang *et al*., 2024).

### 4.7 Quercetin and rutin treatment

Wild-type **s**eeds were sown directly in soil, and one-week-old seedlings were transplanted into pots. Plants were grown at greenhouse conditions with natural daylight variation, maintaining average day/night temperatures of 28/24°C and relative humidity levels of 60/80%. A concentration of 150 μM quercetin hydrate and 0.5 mM rutin hydrate was sprayed onto six-week-old plants on alternative days as treatments and a 0.1% DMSO treatment was considered as control, followed by 27 days of drought stress (D) and later recovery (R). Physiological measurements were recorded during the drought period using an LI-600 porometer (LI-COR Biosciences, USA).

### 4.8 Total flavonoid content, H_2_O_2_ quantification

Total flavonoid content (TFC) was quantified using the aluminum chloride colorimetric method (Baba and Malik, 2015) and H_2_O_2_ quantification was carried out with the ferrous ammonium sulfate/xylenol orange (FOX) method (Sakil *et al*., 2023). Detailed method in Method S10.

### 4.9 Lignin, cellulose and flavonol staining

Lignin staining performed using Wiesner reagent method (Johansen, 1940) and cellulose staining performed using Calcofluor White dye (Bidhendi *et al*., 2020). Detailed method in Method S11. Flavonol staining performed using DPBA staining method (Nguyen, 2020). Detailed method in Method S12.

### 4.10 Statistical analysis and data visualization

All expression data, phenotypic data, and biochemical assay data were analyzed and visualized using GraphPad Prism v10 (GraphPad Software, San Diego, USA). Statistical significance among genes and treatments was evaluated using two-way analysis of variance (ANOVA), followed by Fisher’s least significant difference (LSD) test for multiple comparisons. Differences among means were denoted using a compact letter display (CLD), where values having the same letter indicate no significant difference at p<0.05.

## Author Contributions. CRediT author statement

**Jismon Jose**: Investigation, Conceptualization, Formal analysis, Methodology, Visualization, Writing - Original Draft, Writing - Review & Editing. **Kalyani Prasad**: Resources, Investigation. **Vismaya C**: Investigation. **Swathi Ghantasala**: Investigation. **Palakolanu Sudhakar Reddy**: Resources, Writing - Review & Editing. **Swarup Roy Choudhury**: Conceptualization, Methodology, Supervision, Funding acquisition, Resources, Writing - Original Draft, Writing - Review & Editing

## Acknowledgements

The authors are grateful for funding by Indian Institute of Science Education and Research (IISER), Tirupati and International Crops Research Institute for the Semi-Arid Tropics (ICRISAT), Hyderabad; JJ and VC acknowledges the Council of Scientific and Industrial Research (CSIR), Government of India for Ph.D. fellowship. SG acknowledges DST for the DST Inspire PhD fellowship. The authors are thankful to the members of the Plant Signaling Lab at IISER Tirupati for their help and support.

## Data Availability Statement

The raw RNA sequencing data have been submitted to the NCBI Sequence Read Archive (SRA) database under the BioProject ID: PRJNA1476535 (BioSample: SAMN60716322-SAMN60716333). Additional data generated in this study are available in the supporting information files.

## Supplementary files

**1. Supplementary figures**
**2. Supporting methods**
**3. Supplementary Tables** **Supplementary Table 1:** Groundnut drought stress datasets used in meta-analysis. **Supplementary Table 2:** Meta-analysis results of early drought stress datasets. Log2FoldChange (LFC) of each comparison with meta-analysis statistics. **Supplementary Table 3:** Meta-analysis results of late drought stress datasets. Log2FoldChange (LFC) of each comparison with meta-analysis statistics. **Supplementary Table 4:** Differentially expressed genes obtained from three comparisons from transcriptome analysis **Supplementary Table 5:** Primers used in this study

