## Supplementary material for "Heterotrimeric Gβ subunit orchestrates flavonoid metabolism governing drought-tolerance in groundnut (*Arachis hypogaea* L.)": Supplementary_Figures.pdf

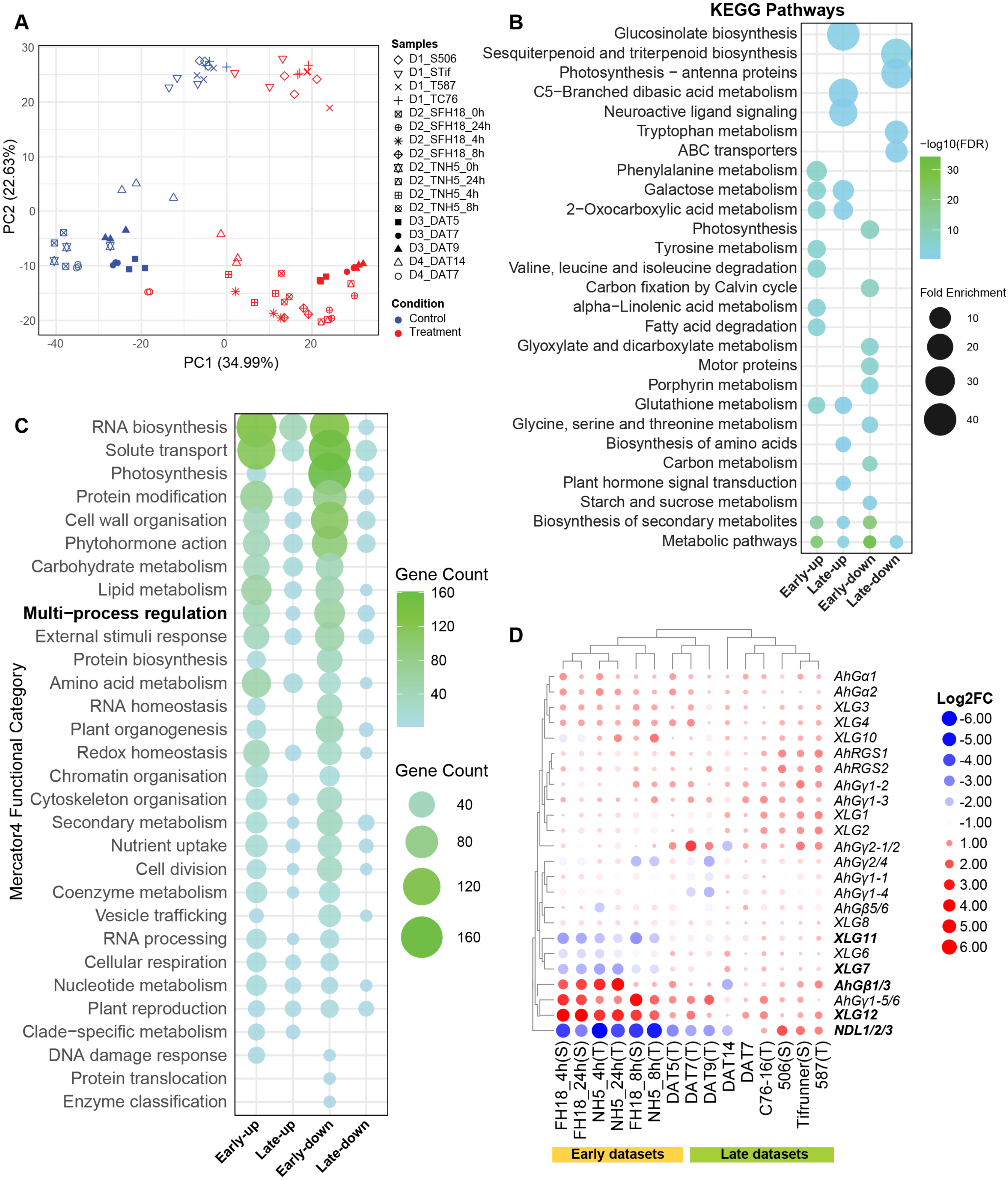


**Supplementary Figure 1. Meta-transcriptome analysis of groundnut drought datasets.** (A) The PCA plot represents the control and drought samples (17) across four datasets. (B) The bubble plot represents the KEGG pathway enrichment across the up- and down-regulated genes from early and late time points. The colour scale represents the significance as -log10(FDR), and the bubble size denotes the fold enrichment. (C) The bubble plot represents the number of genes in different functional categories of Mercator annotation across the up- and down-regulated genes from early and late time points. Both color scale and bubble size represent the gene number. (D) The heatmap displays the log2FC expression of all G-protein components from meta-analysis across early and late time points. Both colour scale and bubble size denote the log2FC values. Significant components are represented in bold font.

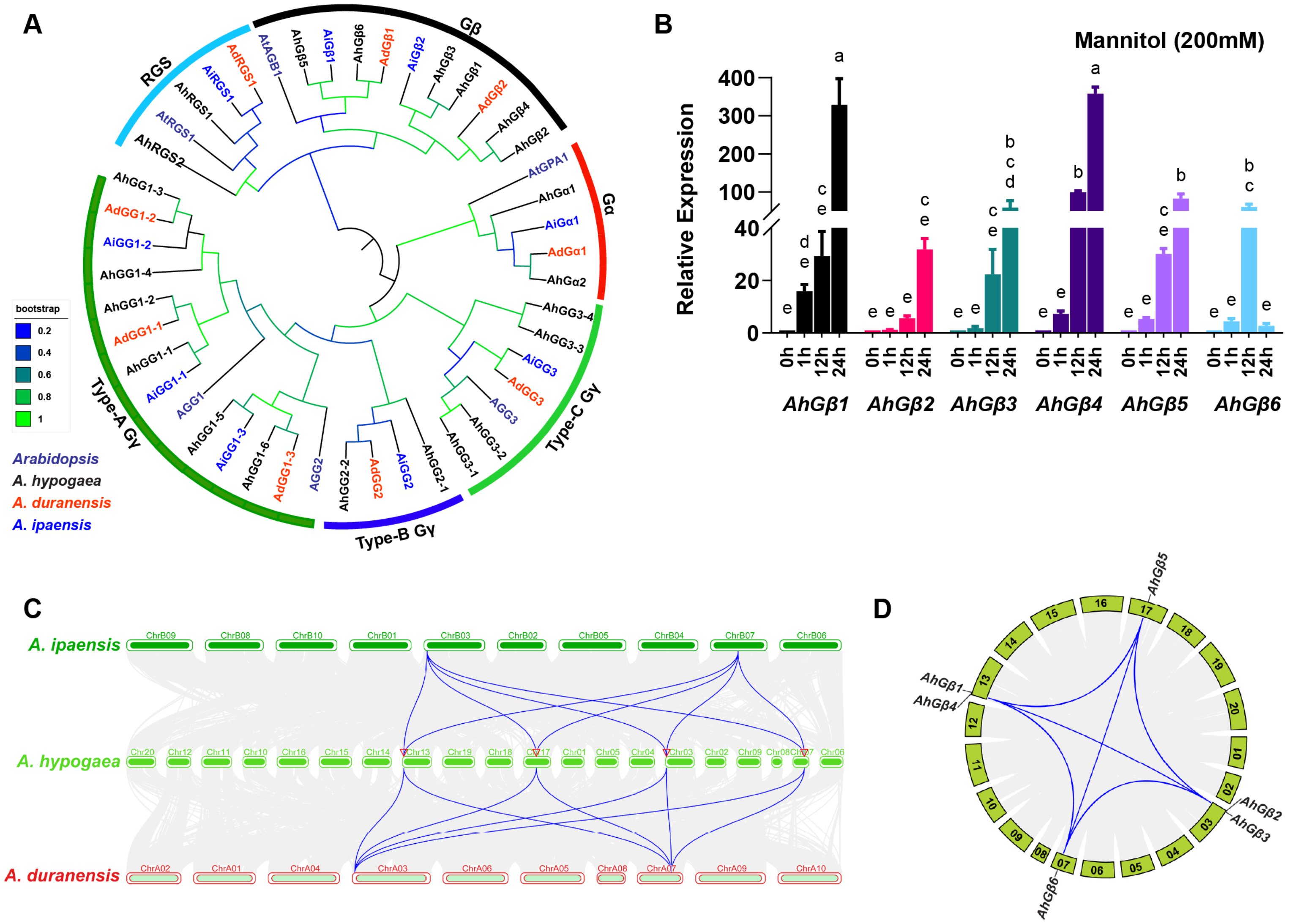

**Supplementary Figure 2. Evolutionary analysis of Gβ subunits in groundnut.** (A) The multi-gene phylogenetic tree exhibits that G-protein subunits (Gα, Gβ, Gγ and RGS) orthologs from *Arachis hypogaea*, *A. duranensis, A. ipaensis*, and *Arabidopsis thaliana*. The branch colour corresponding to the bootstrap value and font colour represent the species. (B) The bar graphs display the relative expression of all six *AhGβ* genes at different time points after treatment with 200 mM mannitol. The letters on top of bars indicate the statistically significant differences ($p<0.05$, two-way ANOVA with Fisher's LSD test). (C) The collinearity syntenic plot between *A. hypogaea*, with its diploid progenitors, *A. duranensis* (A genome), and *A. ipaensis* (B genome) shows the evolutionary link of *Gβ* genes with their parental genomes. (D) The circos plot shows the syntenic region within the *A. hypogaea* genome, and the links indicate conserved synteny among *Gβ* genes.

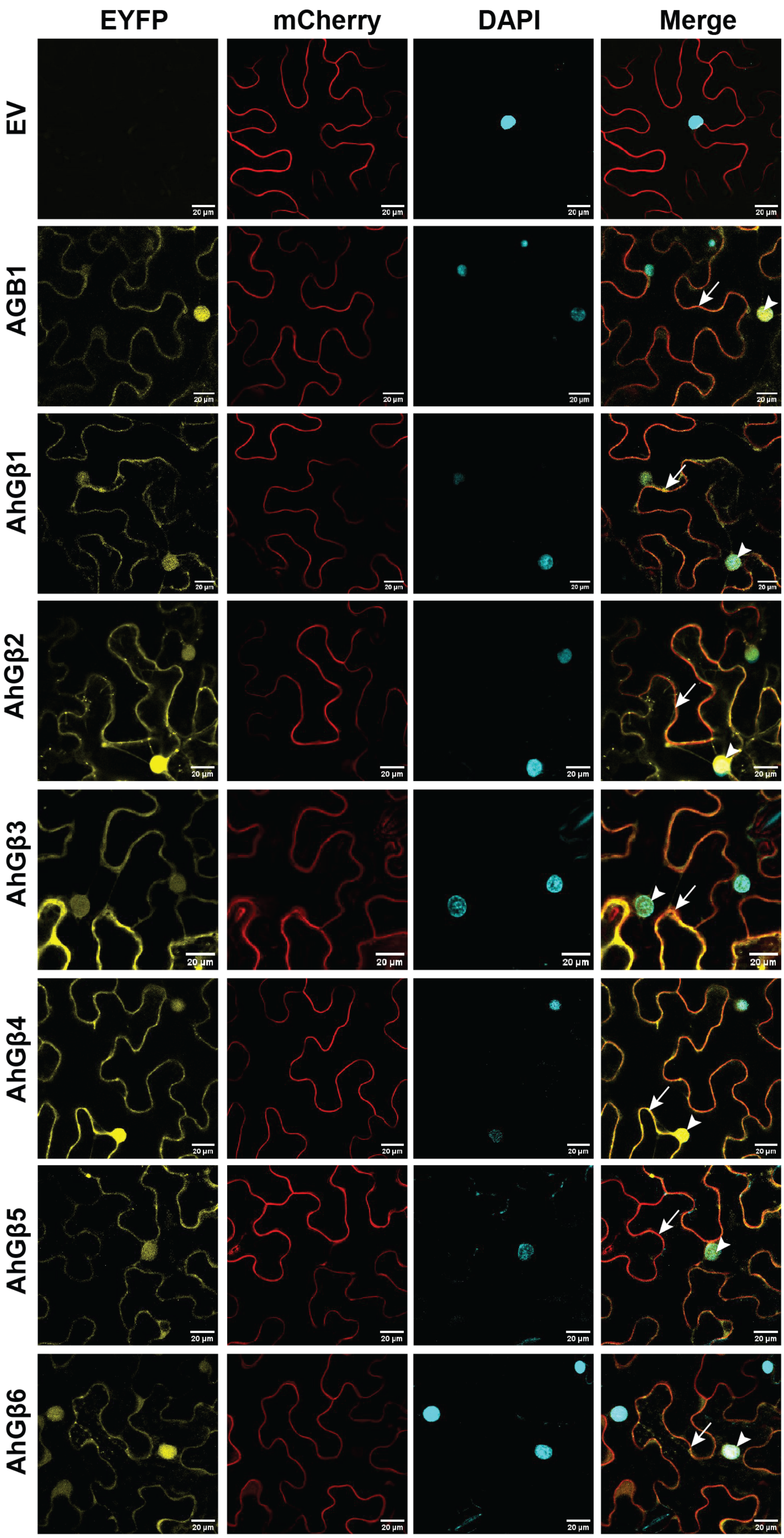


**Supplementary Figure 3. Subcellular localization of AhGβ subunits in *Nicotiana benthamiana*.** Localization of AhGβ subunits with Arabidopsis homolog AGB1. The EYFP panels represent the YFP signal from AhGβ: EYFP proteins, the mCherry panels represent RFP signal from the plasma membrane marker (PIP2: mCherry), the DAPI panels represent the nuclear signal from DAPI channel, and merge panel represent the merged signal from EYFP, mCherry and DAPI channels. EV101 represents the empty vector (pEG101) negative control. The arrow in the merge panel marks plasma membrane localization and the arrow head marks the nuclear localization.

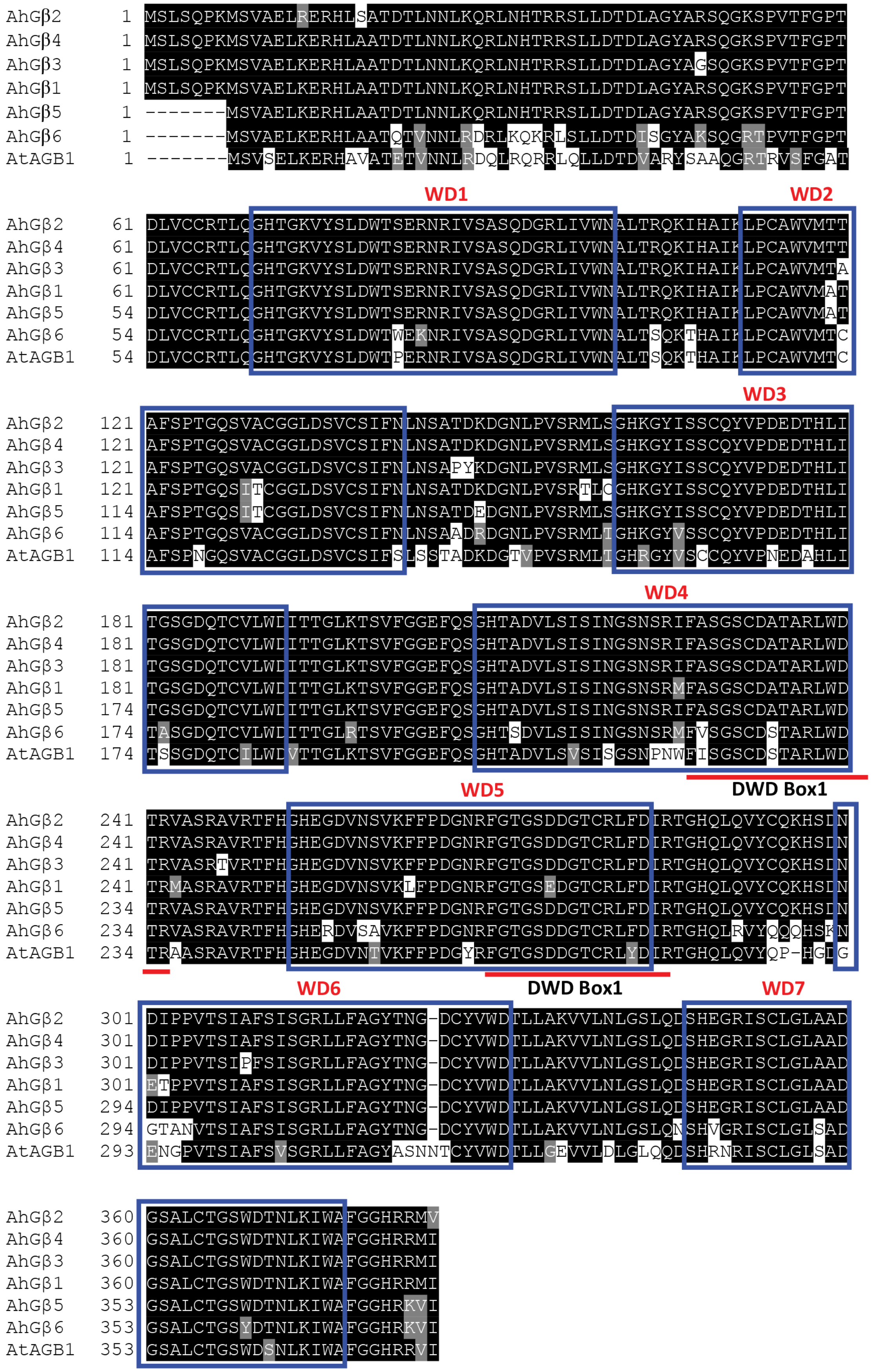


**Supplementary Figure 4. Multiple sequence alignment of AhGβs (1-6) with their Arabidopsis homolog AGB1.** (A) The amino acid sequence alignment of AhGβs proteins from *Arachis hypogaea* (AhGβ1 to AhGβ6) and Arabidopsis (AGB1). The blue square boxes denote the seven WD40 domains [short structural motif of approximately 40 amino acids and often terminating in a tryptophan-aspartic acid (W-D) dipeptide] required for protein-protein or protein-DNA interaction, and the DWD box motifs are predicted to be involved in interaction.

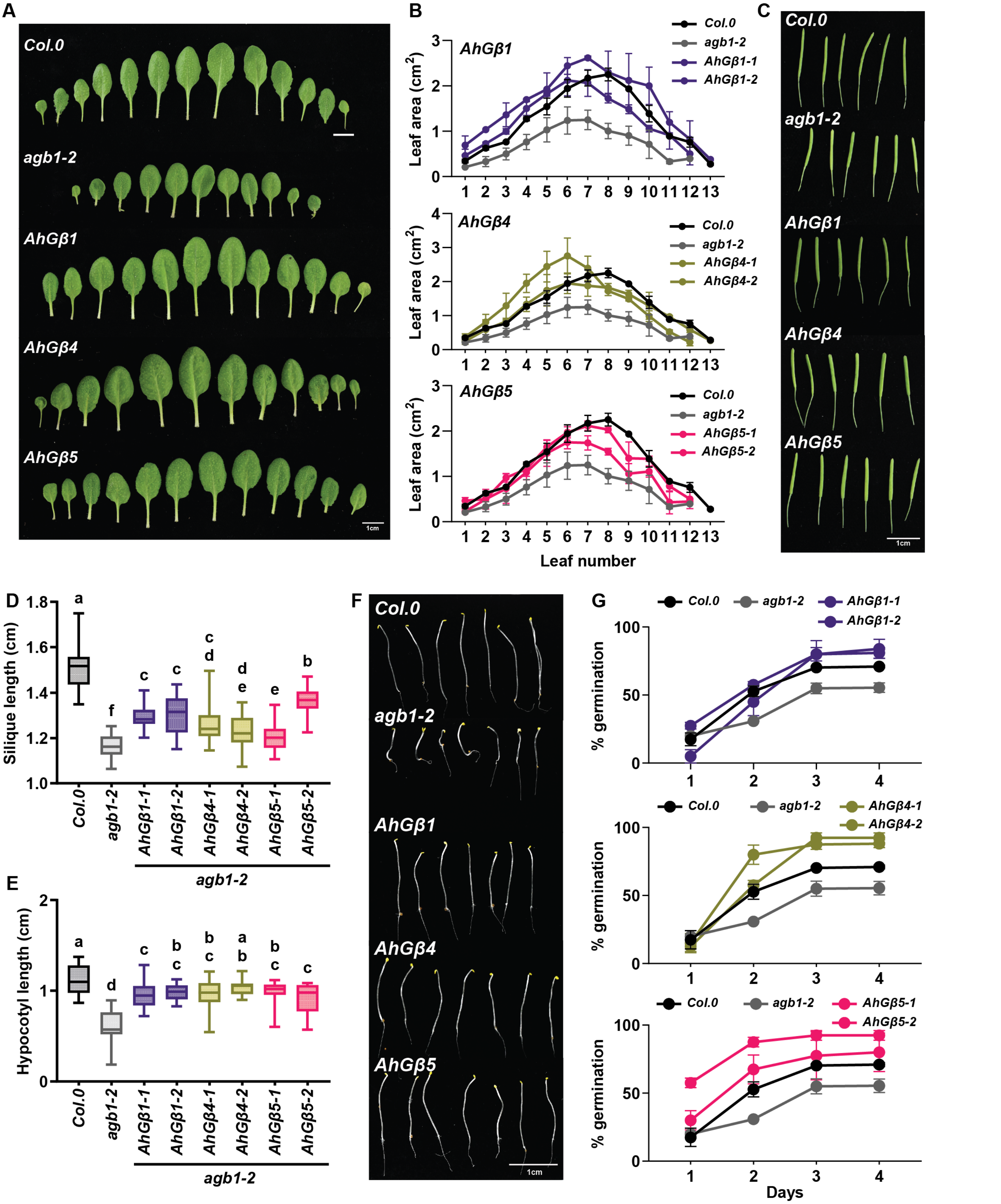


**Supplementary Figure 5. AhGβs complementation studies in agb1-2 mutants.** The morphological features of AhGβ complementation lines in *agb1-2* with mutant (*agb1-2*) and wild-type (*Col.0*) control. (A) Photograph of leaf morphological features, scale bar 1 cm. (B) The line graph represents the leaf area measurement of sequential leaves (from Figure S5A) for *AhGβ1, AhGβ4* and *AhGβ5* complementation lines. (C) Photographs represent the silique morphological features, scale bar 1 cm. (D) The box plot represents the silique length. (E) The box plot represents the hypocotyl length. The middle line in the box plot represents medians, and box range indicates the 25th to 75th percentile. The whiskers denote the minimum-to-maximum range. The letters on top of boxes indicate the statistically significant differences ($p<0.05$, one-way ordinary ANOVA with Fisher's LSD test). (F) Photographs show dark-grown hypocotyl morphological features; scale bar 1 cm. (G) The line graphs represent germination percentage of *AhGβ* complementation lines across 4 days in 3 µM ABA with mutant and wild-type control.

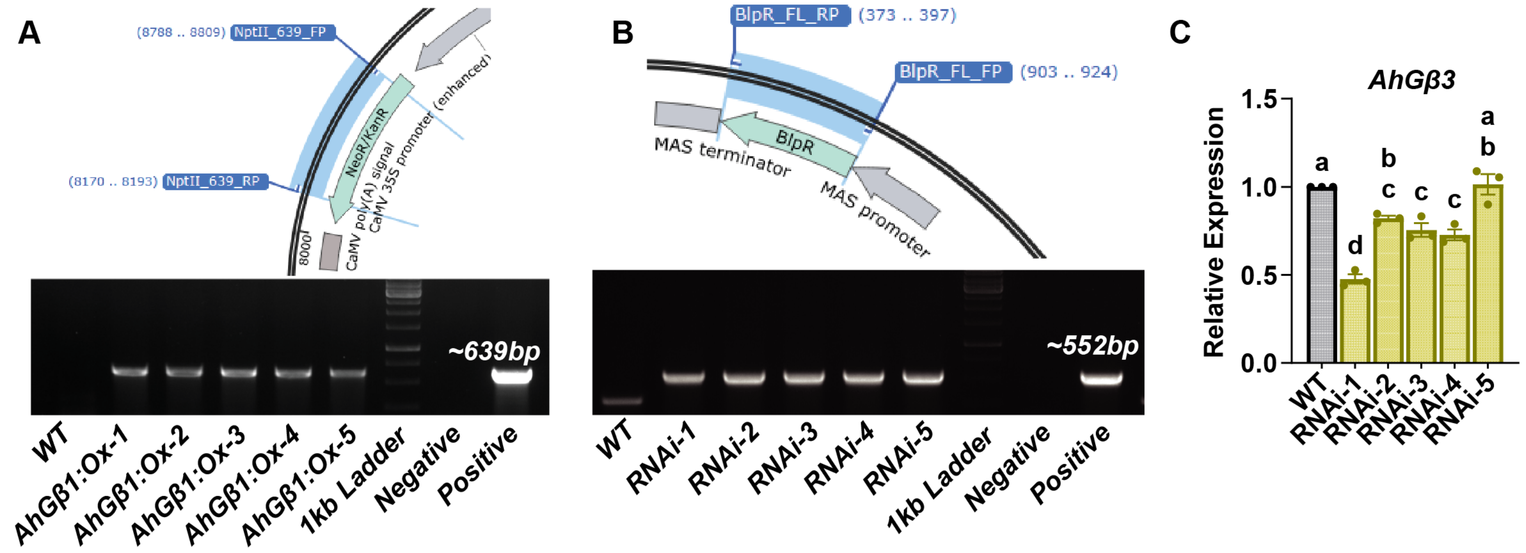


D

| Symbol | Targets (ann1 based) | Total siRNA hits | Efficient siRNA hits |
|---|---|---|---|
| **AhGβ1** | arahy.Tifrunner.gnm1.ann1.VS3LYX.2 | 374 | 21 |
| **AhGβ1** | arahy.Tifrunner.gnm1.ann1.VS3LYX.1 | 374 | 21 |
| **AhGβ3** | arahy.Tifrunner.gnm1.ann1.BV32DL.1 | 301 | 13 |
| AhGβ4 | arahy.Tifrunner.gnm1.ann1.NCS46Q.1 | 180 | 5 |
| AhGβ2 | arahy.Tifrunner.gnm1.ann1.NG5KDE.1 | 165 | 3 |
| AhGβ5 | arahy.Tifrunner.gnm1.ann1.53VILX.2 | 23 | 1 |
| AhGβ5 | arahy.Tifrunner.gnm1.ann1.53VILX.1 | 23 | 1 |
| AhGβ6 | arahy.Tifrunner.gnm1.ann1.SA4SRM.2 | 23 | 1 |
| AhGβ6 | arahy.Tifrunner.gnm1.ann1.SA4SRM.1 | 23 | 1 |

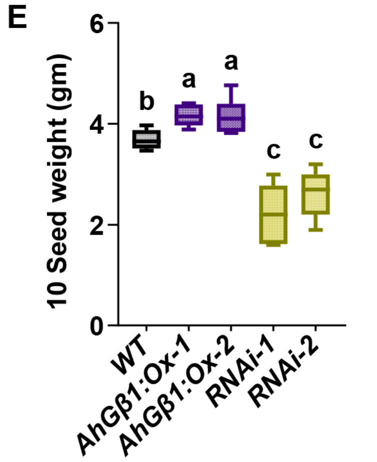


**Supplementary Figure 6. AhGβ1 overexpression and RNAi knock-down in groundnut.** (A) The graphical representation of the *NptII* gene region (pCAMBIA2301) was used to genotype overexpression lines. The gel images show the amplification of *NptII* gene from putative transformants of *AhGβ1:Ox* lines with wild-type (WT), positive control, and negative controls (non-template). (B) The graphical representation of the *Bar* gene region (pFGC5941) is used to genotype the knockdown line. The gel image shows amplification of the *Bar* gene from putative transformants of *RNAi*-knockdown lines, with wild-type (WT), positive control, and negative control (non-template). (C) The bar graphs represent the relative expression of the *AhGβ3* gene in the RNAi knockdown lines compared to the WT. (D) The table represents the targets and off-targets mRNA identified in the groundnut genome using SciFi21 software from the trigger sequence used for knock-down studies. (E) Box plots show 10 seed weights across *WT*, *AhGβ1:Ox*, and *RNAi* lines in control conditions. The middle line in the box plot represents the median, and the box range indicates the 25th to 75th percentiles. The whiskers denote the minimum-to-maximum range. The letters on top of boxes/bars indicate the statistically significant differences ($p<0.05$, one-way ordinary ANOVA with Fisher's LSD test).

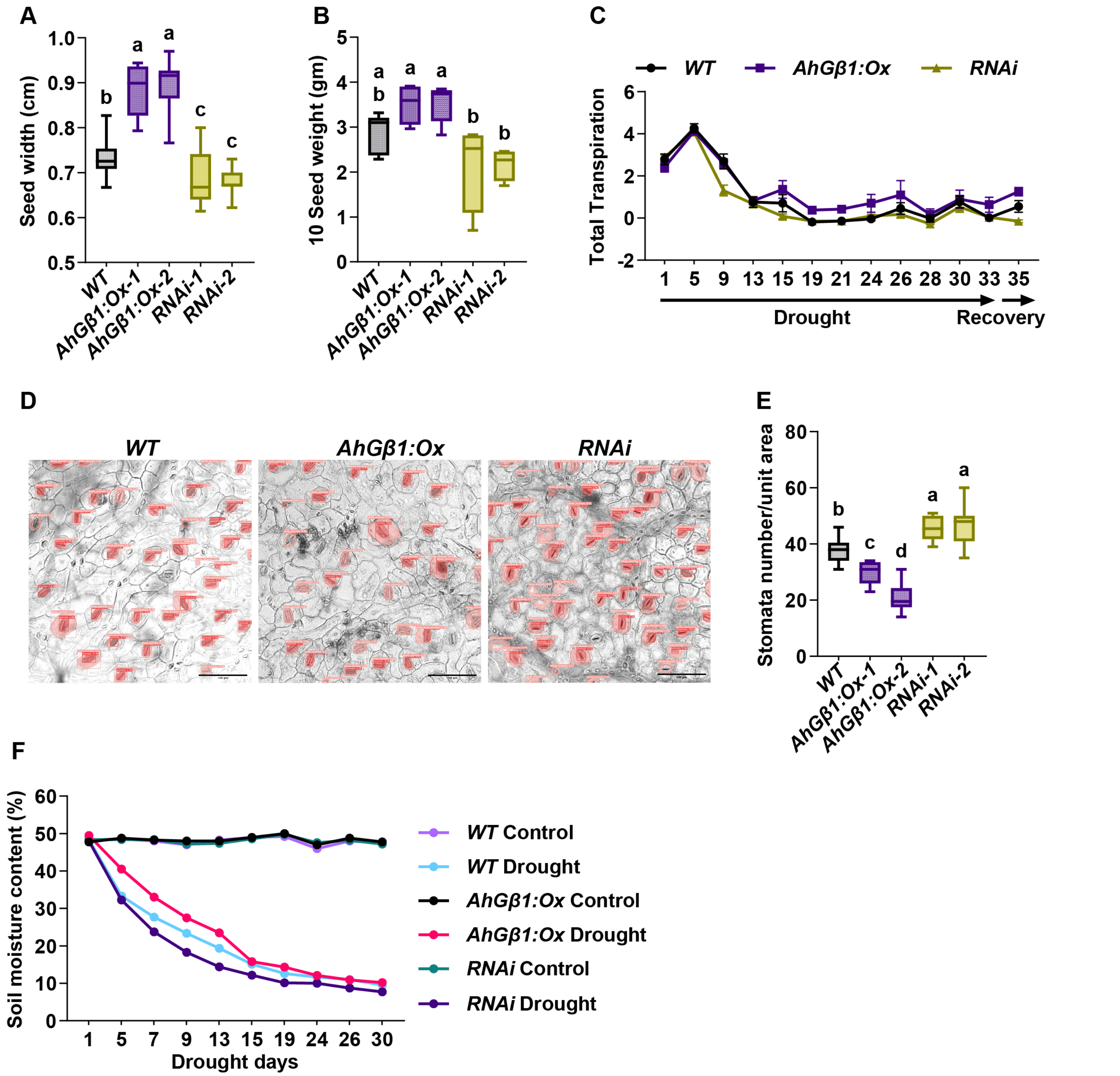


**Supplementary Figure 7. AhGβ1 overexpression and RNAi knockdown in groundnut drought-tolerance**. (A-B) Box plots show seed width (A) and 10 seed weight (B) across *WT, AhGβ1:Ox*, and *RNAi* lines in drought conditions. (C) The line graphs show the total transpiration of *WT, AhGβ1:Ox*, and *RNAi* lines across the drought and recovery phases. (D) The images exhibit the stomatal distribution on the abaxial side of *WT, AhGβ1:Ox*, and *RNAi* lines; scale bar: 100 µm. (E) Box plot represents the stomata number/unit area across *WT, AhGβ1:Ox* and *RNAi* lines. (F) The line graphs show the relative soil moisture content of *WT, AhGβ1:Ox*, and *RNAi* lines under control and drought conditions over the days of drought treatment. The middle line in the box plot represents the median, and the box range indicates the 25th to 75th percentiles. The whiskers denote the minimum-to-maximum range. The letters on top of boxes indicate the statistically significant differences ($p<0.05$, one-way ordinary ANOVA with Fisher's LSD test).

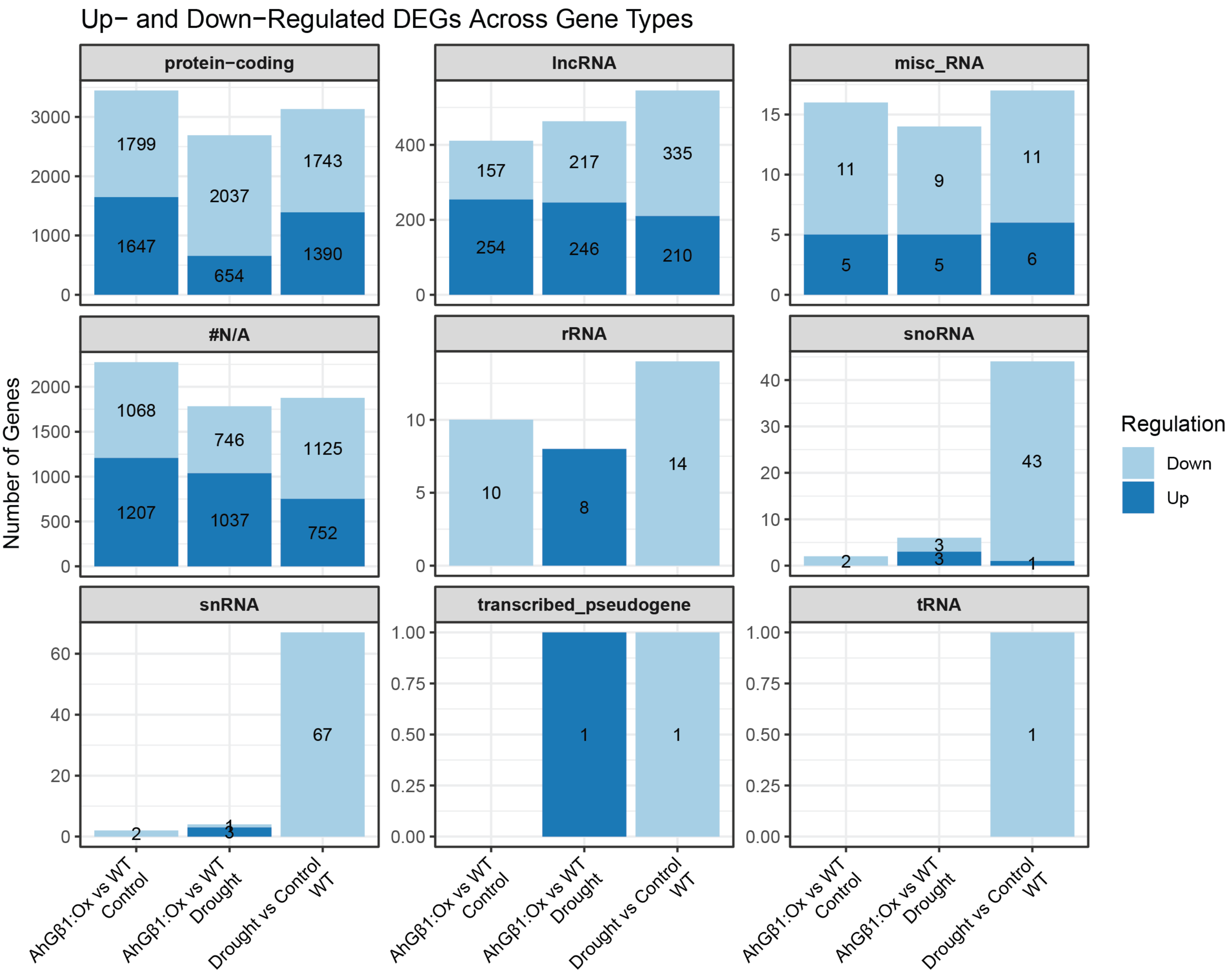


| Comparison | Total_Genes | Total_DEGs | Up | Down |
|---|---|---|---|---|
| AhGβ1:Ox vs WT (Drought) | 71173 | 4970 | 1957 | 3013 |
| AhGβ1:Ox vs WT (Control) | 77109 | 6162 | 3113 | 3049 |
| Drought vs Control (WT) | 73649 | 5699 | 2359 | 3340 |

**Supplementary Figure 8. Drought stress transcriptome analysis of AhGβ1 overexpression lines.** The bar graphs show the distribution of both up- and downregulated genes across different gene types from all three DEG analyses. The bottom table represents the overall details of the three differential comparisons. Dark blue represents the number of upregulated genes, and light blue represents the number of downregulated genes. The X-axis represents the comparisons. Only protein-coding gene type used in the downstream analysis.

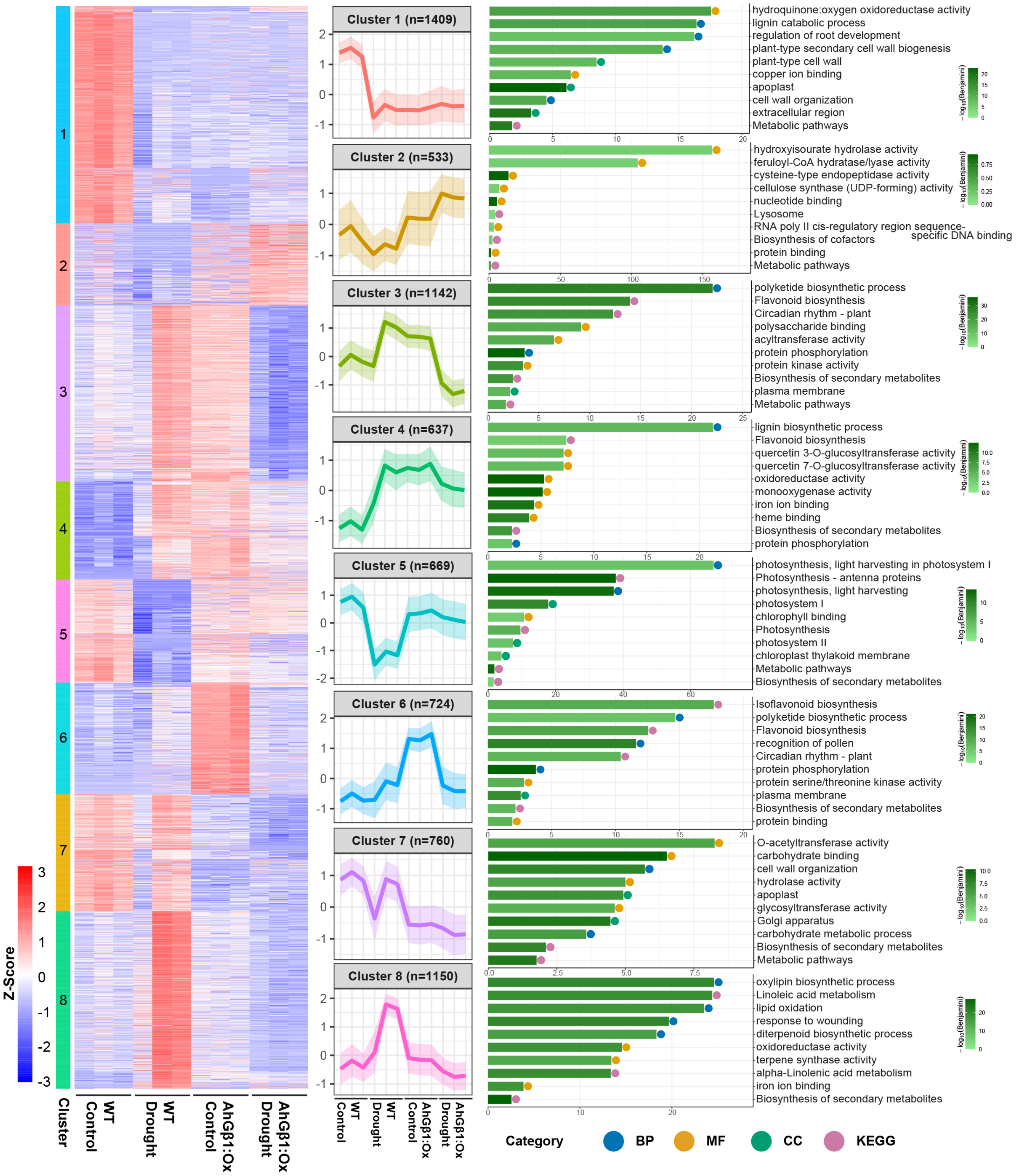


**Supplementary Figure 9. AhGβ1 influences secondary metabolism, photosynthesis, and cell-wall remodeling during drought.** The heatmap shows k-means clustering and z-score expression of total DEGs across the samples. The line graphs represent the average expression curves of each gene cluster across samples (X-axis), with the number of genes in each cluster. The bar graphs show the top 10 GO terms or KEGG pathways for the corresponding cluster. Bubble color represents the category of the term/pathway (BP-biological process, MF-molecular function, CC-cellular component, and KEGG pathway). The bar colour represents the significance as −log10(Benjamini). Three clusters from this figure have been represented in Figure 5C.

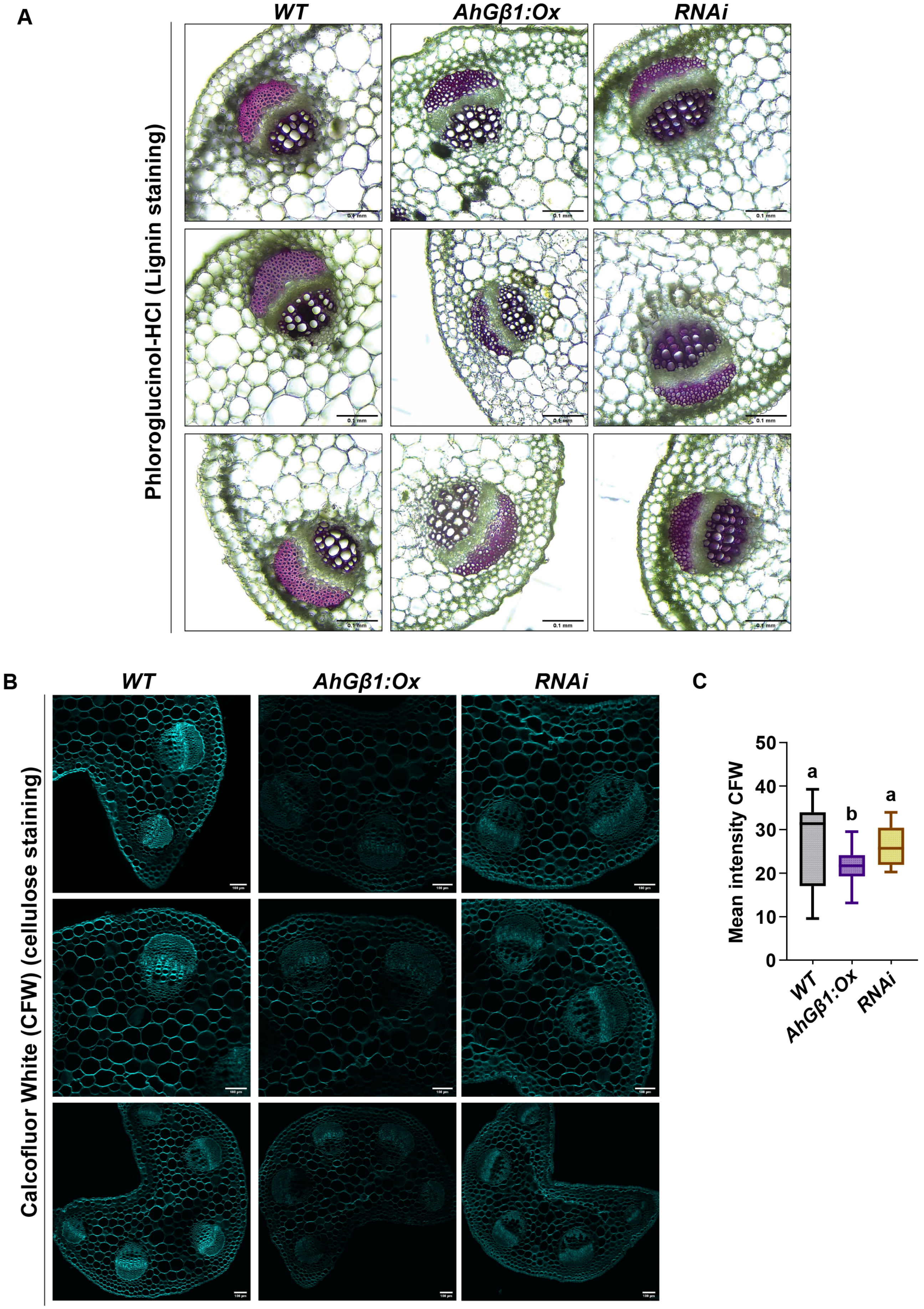


**Supplementary Figure 10. Lignin and cellulose staining.** (A) The images represent the leaf petiole vascular bundles of *WT, AhGβ1:Ox* and *RNAi* lines stained with Wiesner reagent, scale bar 0.1mm. (B) The images presenting the CFW-stained petiole sections of *WT, AhGβ1:Ox*, and *RNAi* lines, scale bar 100 µm. (C) Box plots represent the mean intensity of CFW across *WT, AhGβ1:Ox,* and *RNAi* lines petiole sections. The middle line in the box plot represents the median, and the box range indicates the 25th to 75th percentiles. The whiskers denote the minimum-to-maximum range. The letters on top of boxes indicate the statistically significant differences ($p<0.05$, one-way ordinary ANOVA with Fisher's LSD test).

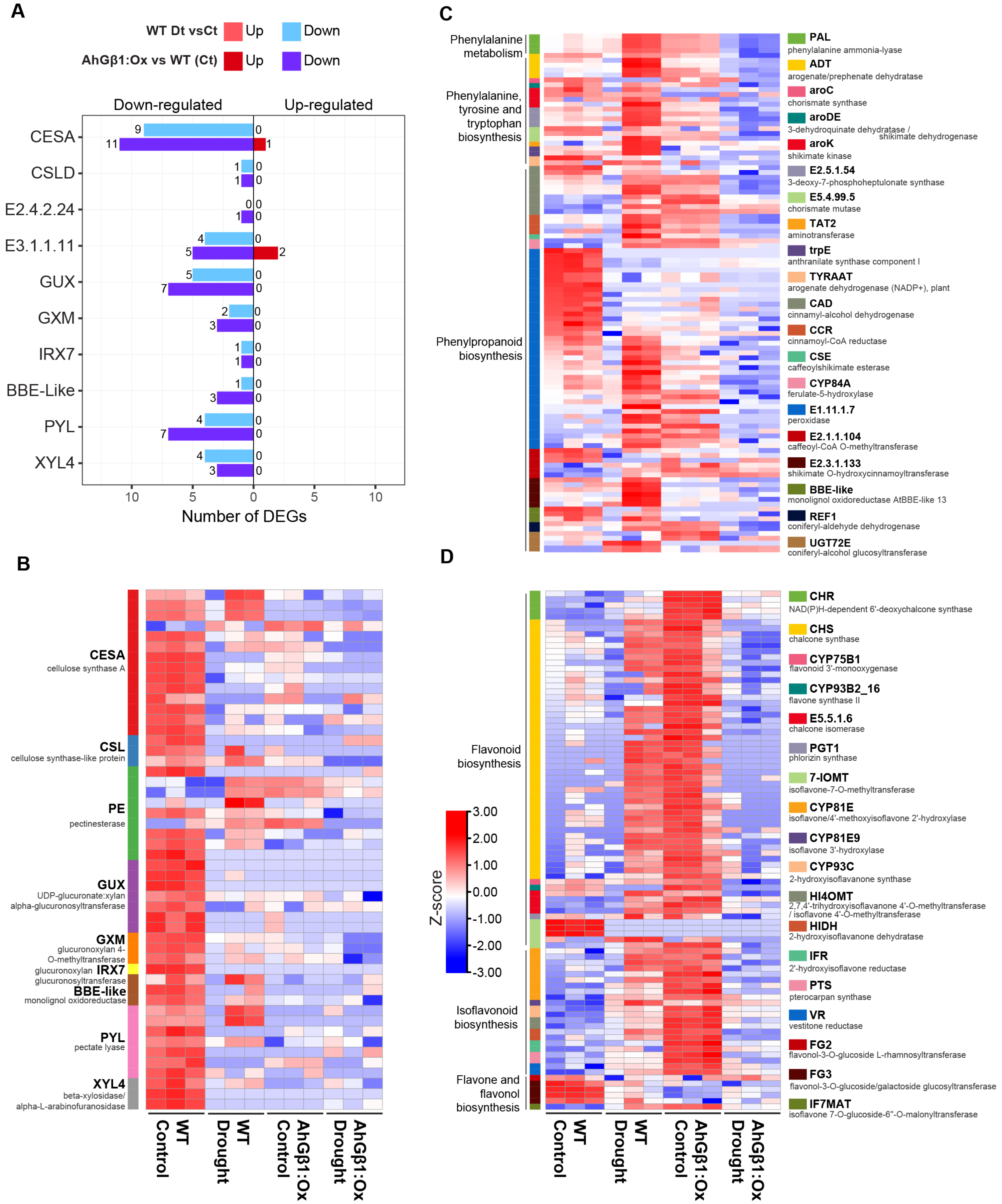


**Supplementary Figure 11. AhGβ1 influences secondary metabolism and cell-wall remodeling during drought.** (A) The mirror bar graphs show the number of DEGs in the different cell wall-related enzyme families for two different comparisons. Color represents up- and down-regulated genes of the corresponding comparison. (B) The heatmap shows the z-score expression of different cell wall-related enzymes, DEGs across samples. (C) The heatmap represents z-score expression of phenylpropanoid biosynthesis-related enzyme DEGs across samples. (D) The heatmap shows the z-score expression of flavonoid-, isoflavonoid-, flavone-, and flavonol biosynthesis-related enzyme DEGs across samples. The colour bar denotes the biosynthetic enzyme groups.

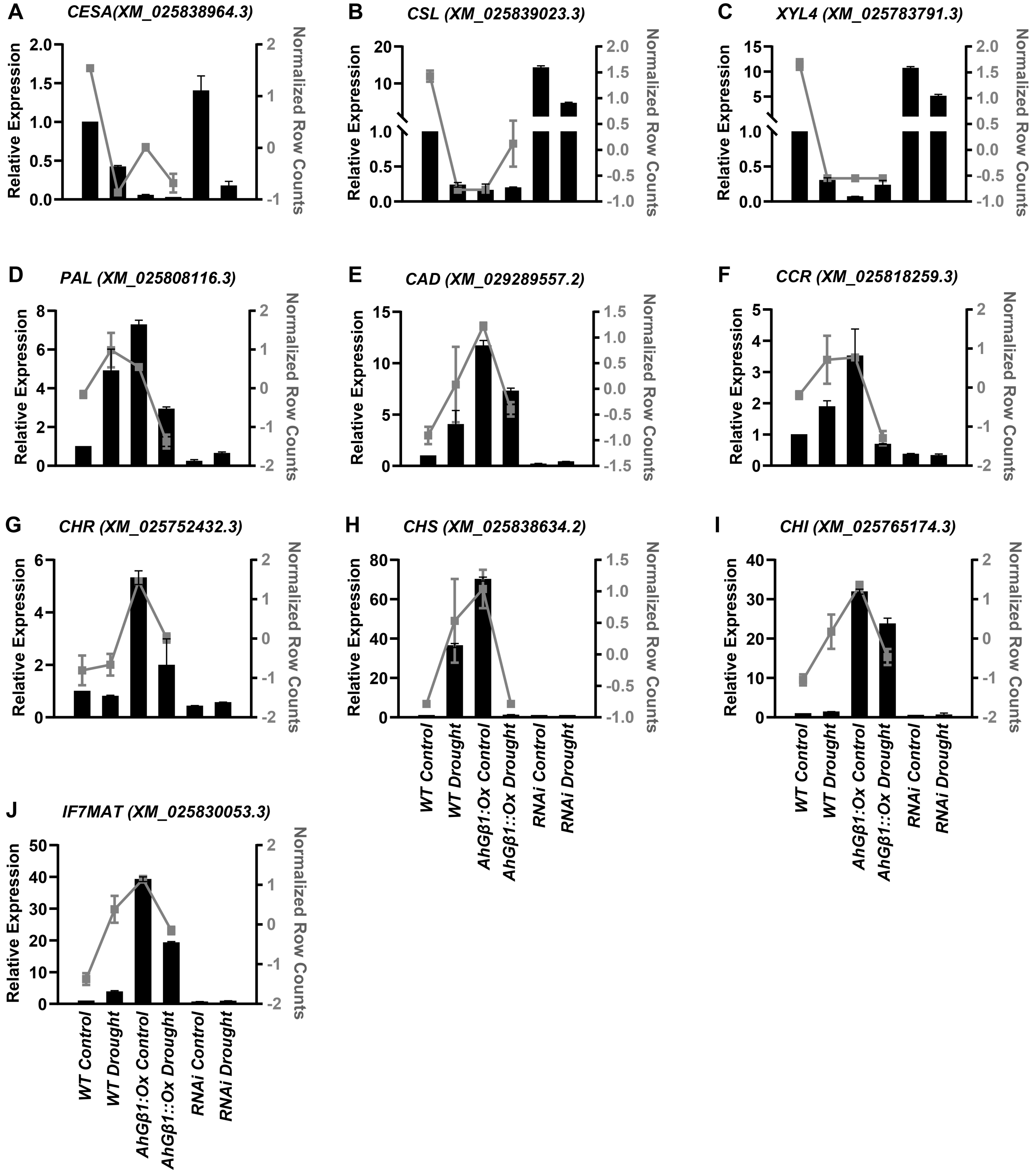


**Supplementary Figure 12. The expression analysis of cell-wall remodeling and secondary metabolism-related genes.** The expression validation of selected genes from transcriptome data. Bar graphs denote qRT-PCR relative expression (left Y-axis) of genes across genotypes and conditions. The line graph represents the normalized row counts (right Y-axis) from transcriptome data across *WT* and *AhGβ1:Ox* under drought and control. All samples are on the X-axis. The selected candidate genes include cell wall-related genes - (A) Cellulose synthase A (*CESA*), (B) CESA-like (*CSL*), and (C) Beta-xylosidase (*XYL4*). Phenylpropanoid pathway related genes - (D) Phenylalanine ammonia-lyase (*PAL*), (E) Cinnamyl alcohol dehydrogenase (*CAD*), and (F) Cinnamoyl-CoA reductase (*CCR*). Flavonoid biosynthesis related genes - (G) Chalcone reductase (*CHR*), (H) Chalcone synthase (*CHS*), (I) and Chalcone isomerase (*CHI)*. Isoflavonoid biosynthesis related gene - (J) Isoflavone 7-O-glucoside-6''-O-malonyltransferase (*IF7MAT*). The NCBI mRNA accession of the corresponding gene is denoted in brackets.

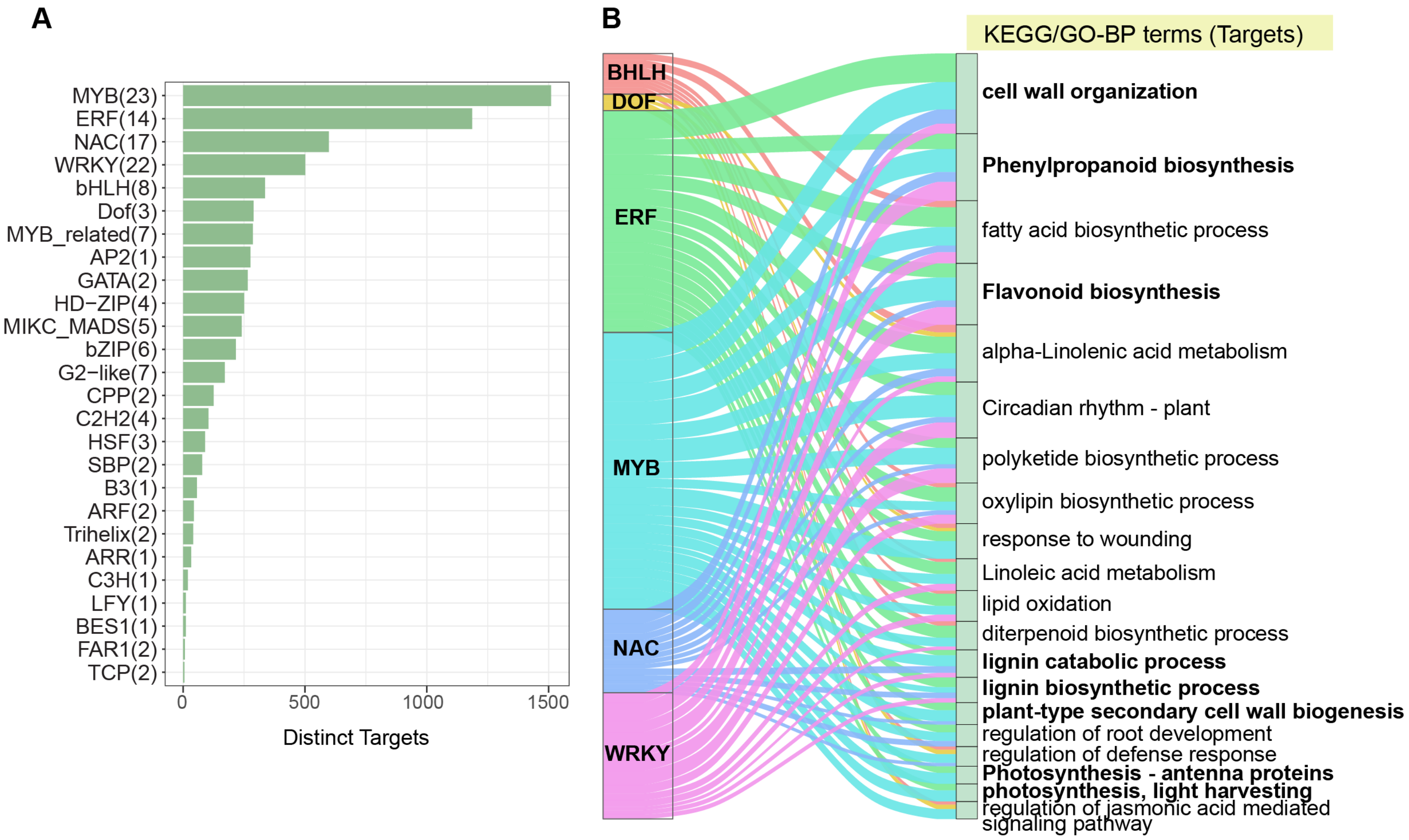


**Supplementary Figure 13. AhGβ1 transcriptionally influences secondary metabolism and cell-wall remodeling during drought.** (A) Bar graphs represent the number of distinct targets (X-axis) across TF families (Y-axis), and the number at the start of the bar denotes the number of TFs in the corresponding family from the DEGs. (B) The alluvial plot is showing the major TF families and their targets' functional pathways/terms (GO/KEGG). Pathways with bold font represent secondary metabolism, cell wall biogenesis, and photosynthesis-related pathways/terms.

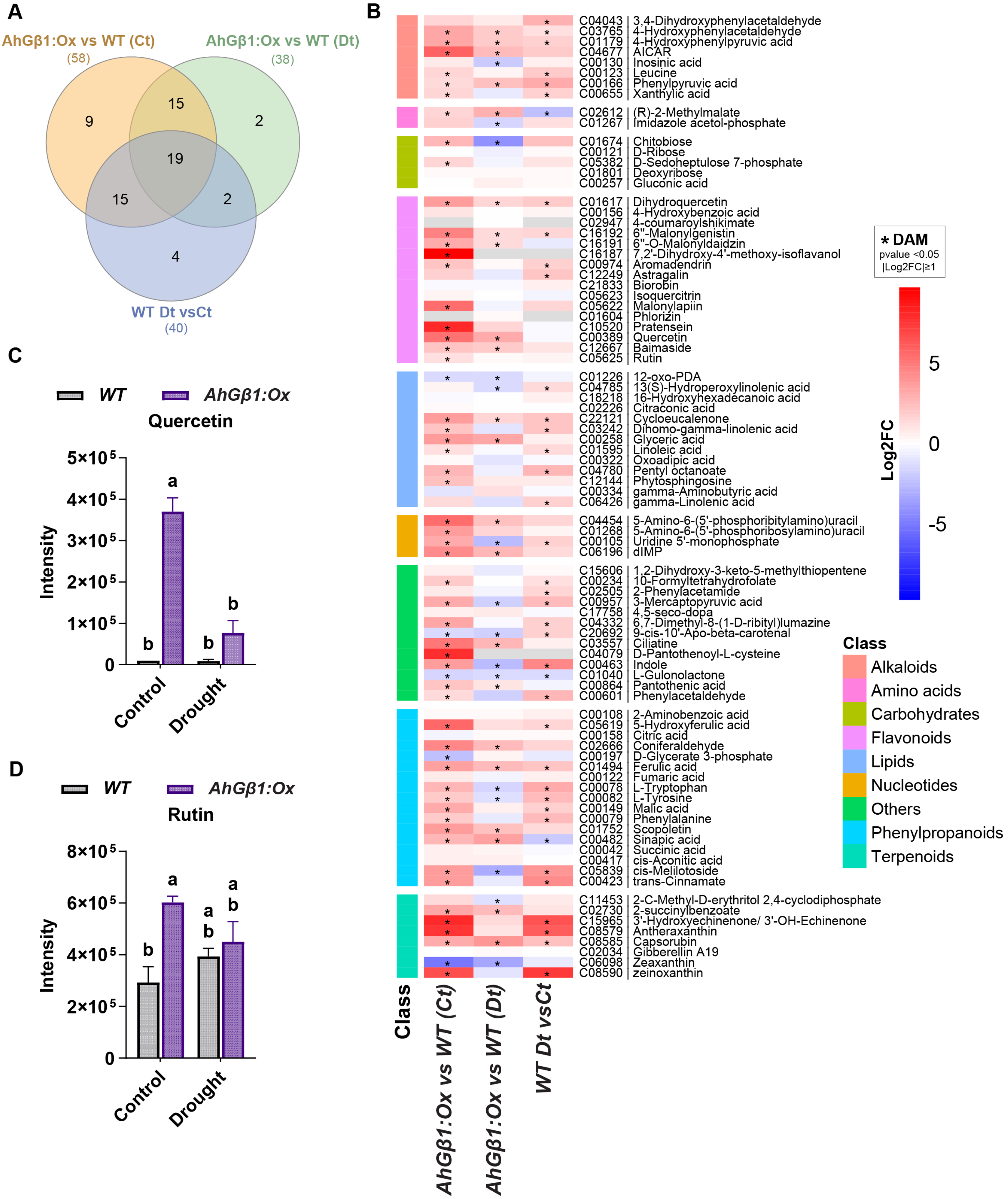


**Supplementary Figure 14. Drought stress metabolomics analysis of AhGβ1 overexpression lines.** (A) The Venn diagram shows the overlapping DAMs (Differentially abundant metabolites) across the comparisons, and the bracket number represents the number of DAMs in the corresponding comparison. (B) The heatmap shows the log2-fold change in metabolite abundance across the three comparisons; the color bar denotes metabolite classes based on the biosynthetic pathways they are involved in; and asterisks indicate significant metabolites. (C-D) The bar graph shows the raw intensity values for quercetin (C) and rutin (D) across the control and drought samples from *WT* and *AhGβ1:Ox* lines. The letters on top of bars indicate the statistically significant differences ($p<0.05$, one-way ordinary ANOVA with Fisher's LSD test).

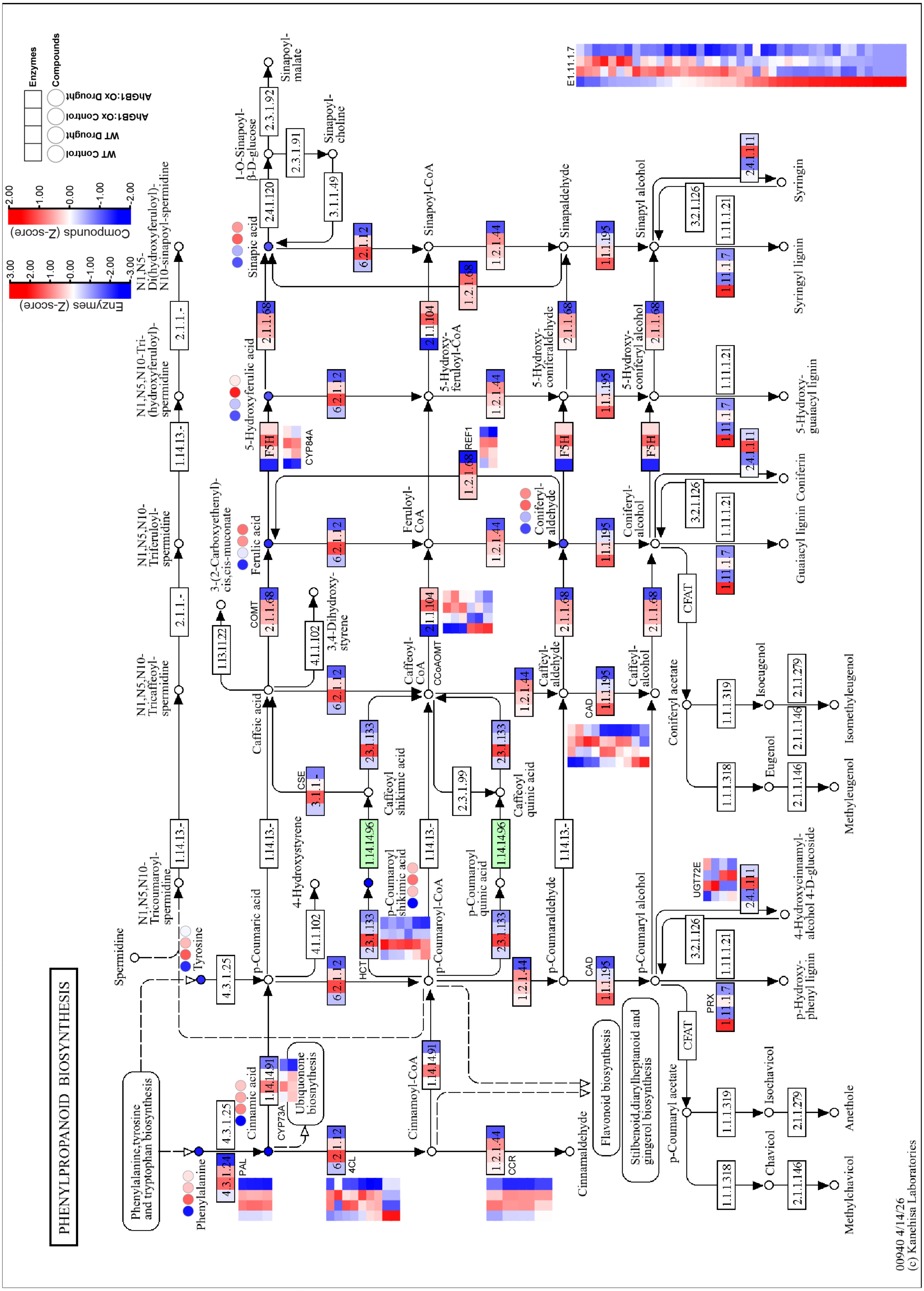


**Supplementary Figure 15. Integration of transcriptome and metabolome: Phenylpropanoid biosynthesis.** The KEGG pathway map of phenylpropanoid biosynthesis represents the z-score expression of biosynthetic enzymes from transcriptome and the mean z-score abundance of compounds from metabolomics analysis across the control and drought samples of *WT* and *AhGβ1:Ox* lines. The heatmap strips (rectangles) represent biosynthetic enzyme expression. Circle heatmap strips represent compound abundance. The colour legends (blue to red) denote the z-score expression or abundance scale of enzymes or compounds.

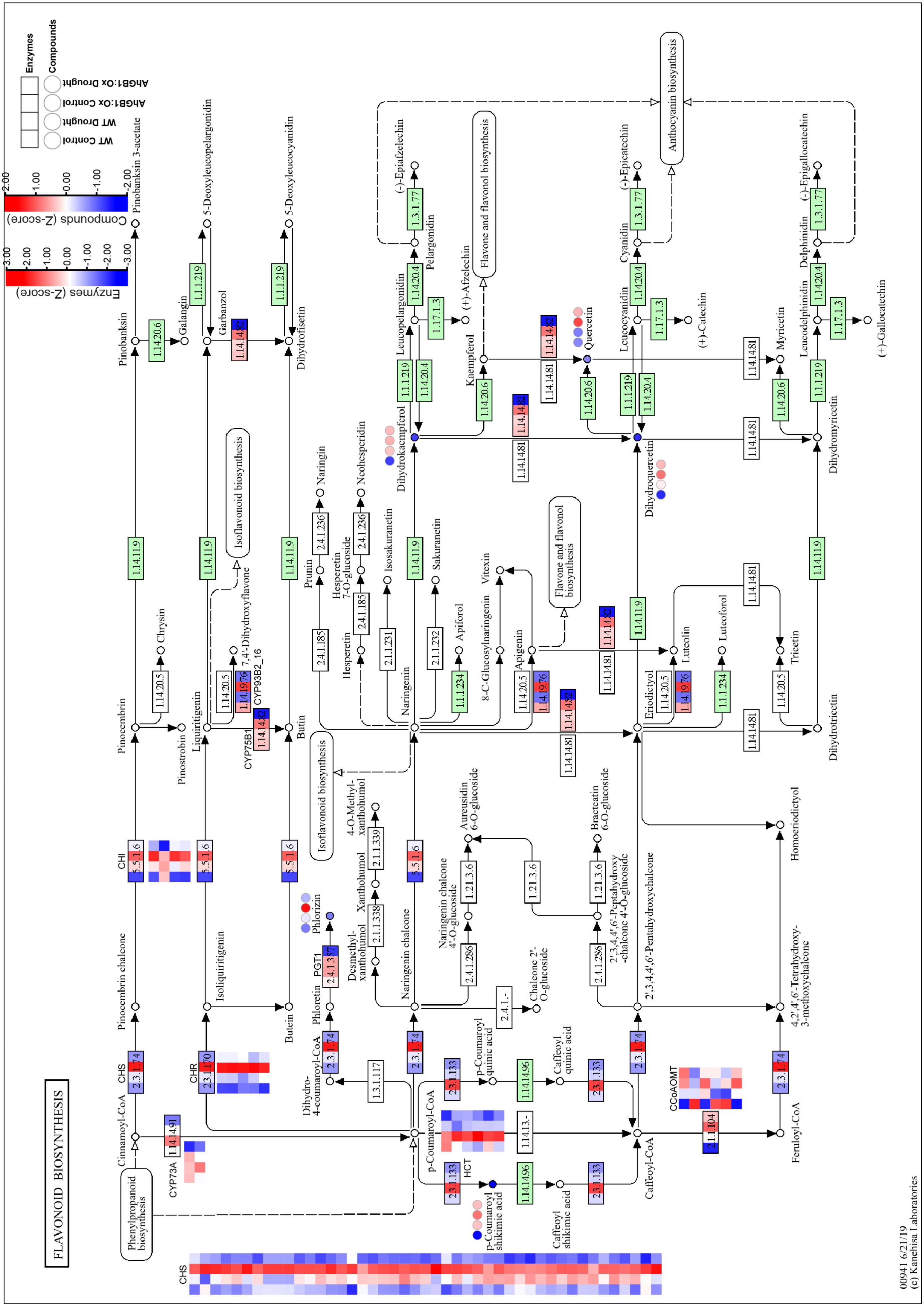


**Supplementary Figure 16. Integration of transcriptome and metabolome: Flavonoid biosynthesis.** The KEGG pathway map of flavonoid biosynthesis represents the z-score expression of biosynthetic enzymes from transcriptome and the mean z-score abundance of compounds from metabolomics analysis across the control and drought samples of *WT* and *AhGβ1:Ox* lines. The heatmap strips (rectangles) represent biosynthetic enzyme expression. Circle heatmap strips represent compound abundance. The colour legends (blue to red) denote the z-score expression or abundance scale of enzymes or compounds.

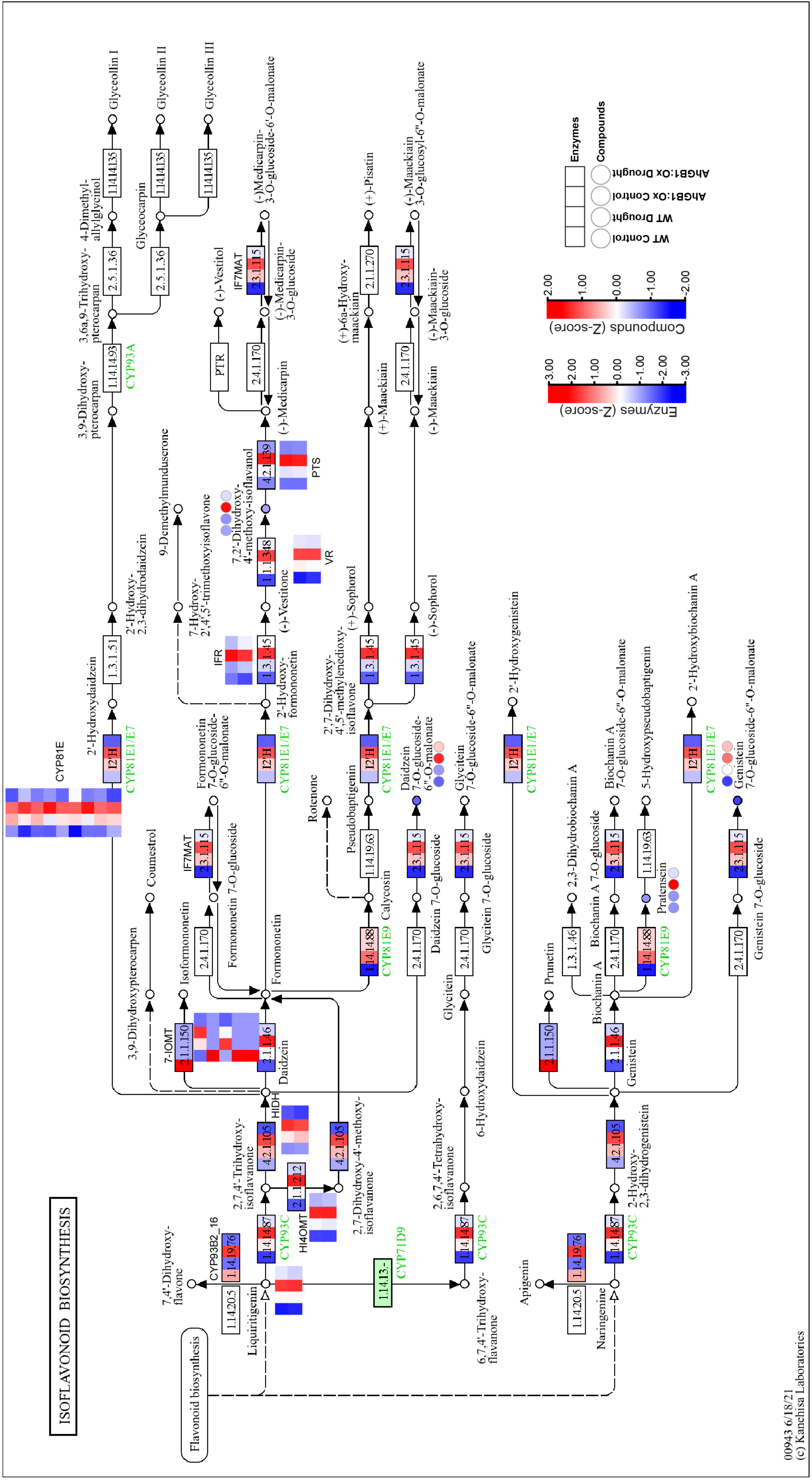


**Supplementary Figure 17. Integration of transcriptome and metabolome: Isoflavonoid biosynthesis.** The KEGG pathway map of isoflavonoid biosynthesis represents the z-score expression of biosynthetic enzymes from transcriptome and the mean z-score abundance of compounds from metabolomics analysis across the control and drought samples of *WT* and *AhGβ1:Ox* lines. The heatmap strips (rectangles) represent biosynthetic enzyme expression. Circle heatmap strips represent compound abundance. The colour legends (blue to red) denote the z-score expression or abundance scale of enzymes or compounds.

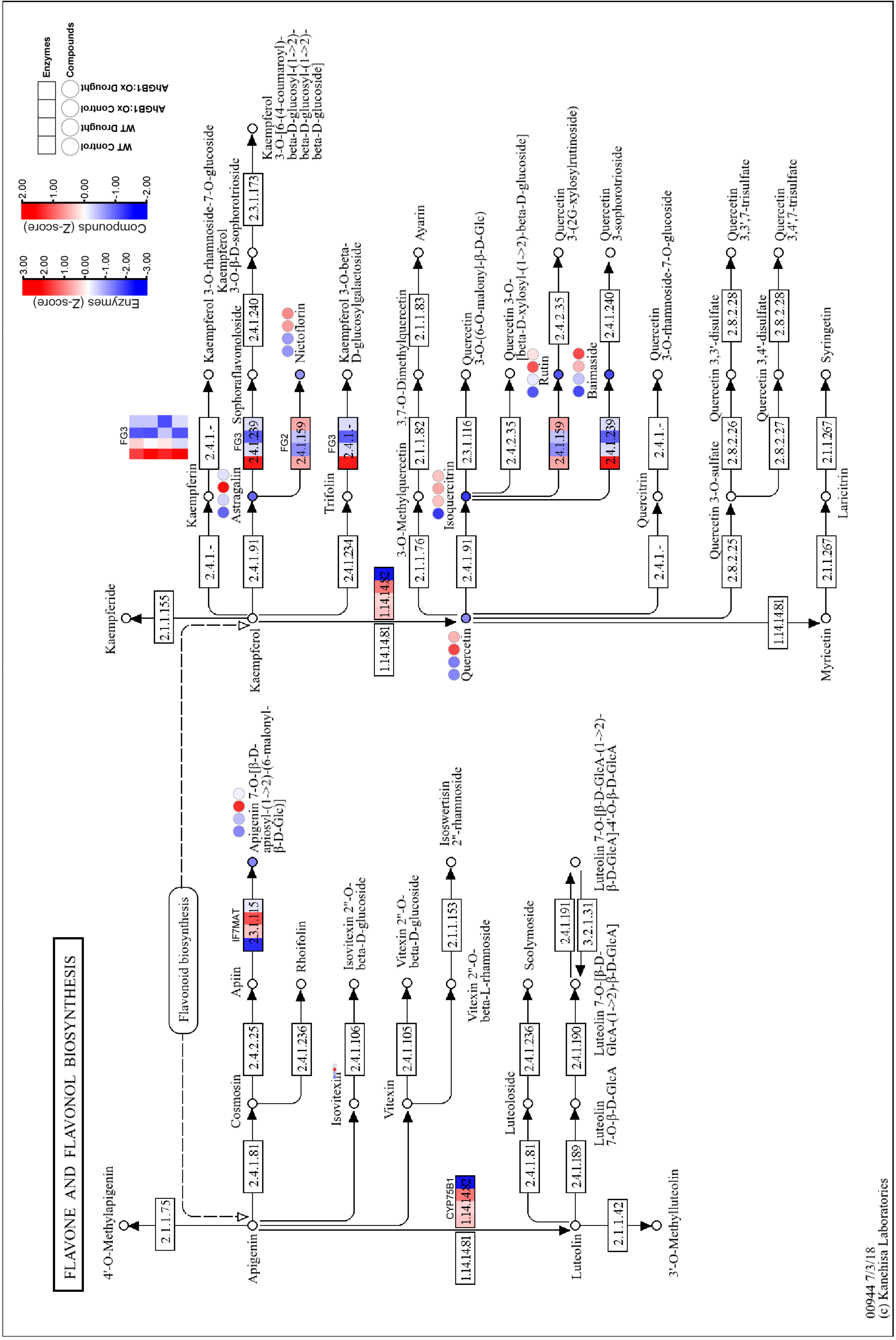


**Supplementary Figure 18. Integration of transcriptome and metabolome: IFlavone and flavonol biosynthesis.** The KEGG pathway map of flavone and flavonol biosynthesis represents the z-score expression of biosynthetic enzymes from transcriptome and the mean z-score abundance of compounds from metabolomics analysis across the control and drought samples of *WT* and *AhGβ1:Ox* lines. The heatmap strips (rectangles) represent biosynthetic enzyme expression. Circle heatmap strips represent compound abundance. The colour legends (blue to red) denote the z-score expression or abundance scale of enzymes or compounds.

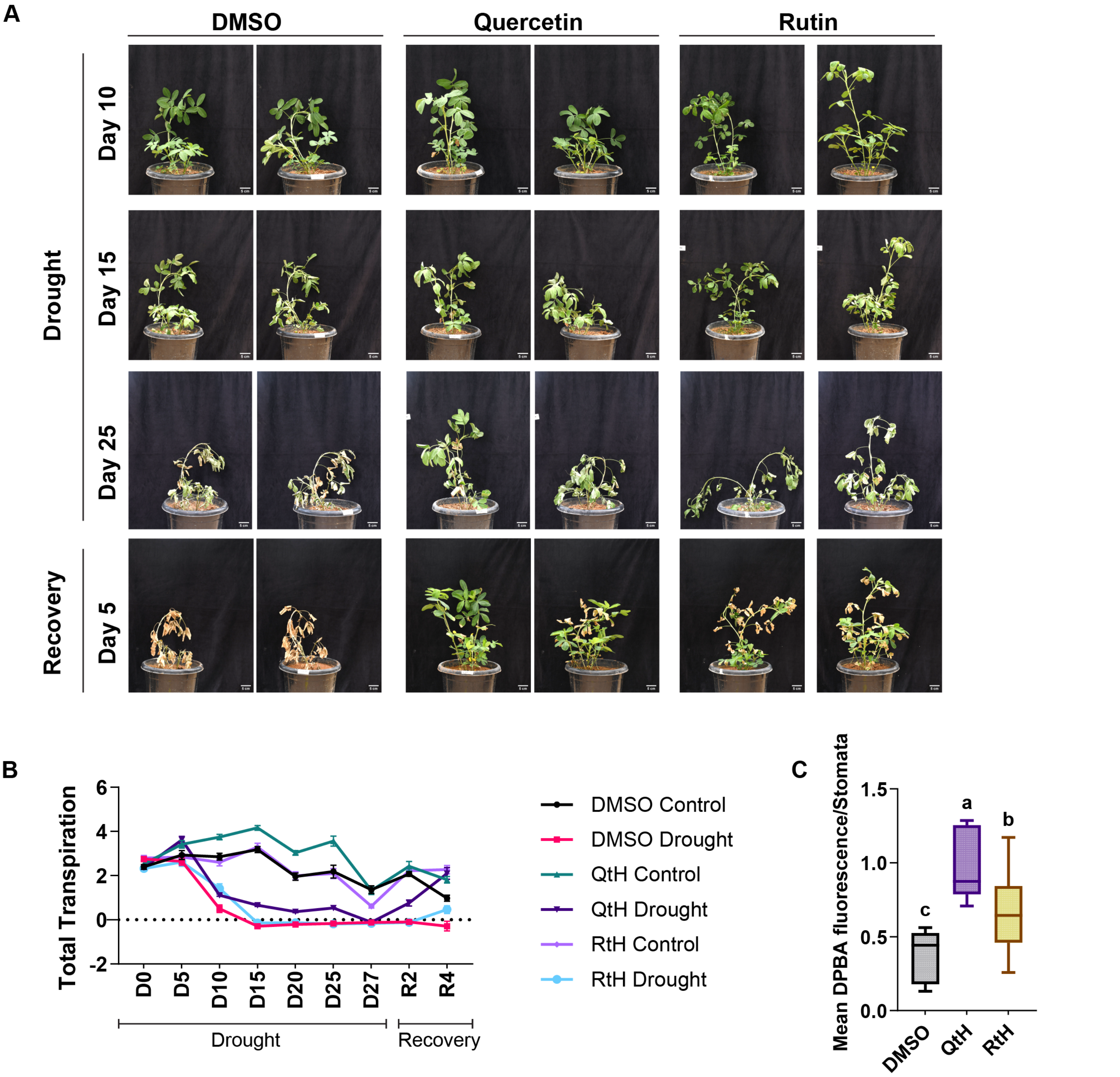


**Supplementary Figure 19. Quercetin and rutin treatments improved drought-tolerance in groundnut.** (A) Photographs show the DMSO-, quercetin-, and rutin-treated plants during the drought, and recovery days. Scale bar 5 cm. 150µM quercetin and 0.5mM rutin applied for three alternate days followed by the 27 days of drought stress (D) and five-day recovery by rewatering (R). DMSO treatment considered as control. (B) Line graphs display the total transpiration across compound treated plants during drought stress (D0 to D27) and recovery (R2 and R4) days. Line color denotes the control and drought plants of DMSO, quercetin (QtH), and rutin (RtH) treatments. (C) Box plots represent the mean intensity of DPBA-flavanol fluorescence per stomata across the DMSO, QtH and RtH treatments. The middle line in the box plot represents the median, and the box range indicates the 25th to 75th percentiles. The whiskers denote the minimum-to-maximum range. The letters on top of boxes indicate the statistically significant differences ($p<0.05$, one-way ordinary ANOVA with Fisher's LSD test).
