## Supplementary material for "Heterotrimeric Gβ subunit orchestrates flavonoid metabolism governing drought-tolerance in groundnut (*Arachis hypogaea* L.)": Supporting_Methods.pdf

### Method S1: Meta-analysis of groundnut drought stress datasets

Initial data quality assessment performed using FastQC (v0.12.1), followed by high-quality reads filtering using Fastp v0.23.4 (Chen *et al.*, 2018). These high-quality reads were aligned to the *Arachis hypogaea* reference genome (Tifrunner.gnm2.J5K5) using HISAT2 (v2.2.1) (Kim *et al.*, 2019) with default parameters. Transcript assembly and quantification were performed from the aligned reads of each sample using StringTie (v2.2.1) (Pertea *et al.*, 2015). The resulting expression read counts for the transcripts were used for differential gene expression analysis with the DESeq2 R package. The comparison was made between treatment and control across 4 datasets, yielding 15 comparisons, which were classified into two groups: early (7) and late (8), based on the time points of drought stress. A meta-analysis was conducted using the metaRNASeq R package (Rau et al., 2014).

### Method S2: Gene expression studies

Total RNA was extracted using the TRIzol reagent (Invitrogen, Cat. No. 15596018) and reverse-transcribed into cDNA using the PrimeScript™ 1st Strand cDNA Synthesis Kit (TaKaRa, Cat. No. 6110B) according to the manufacturer's instructions. Quantitative real-time PCR (RT-qPCR) was performed on a CFX384 Touch Real-Time PCR Detection System (Bio-Rad, Cat. No. 1855484) using gene-specific primers, with *Actin* as the internal reference gene (Supplementary Table 4). Reactions were prepared using TB Green® Premix Ex Taq™ II (Tli RNase H Plus) (TaKaRa, Cat. No. RR820A).

### Method S3: Identification, cloning, and evolutionary analysis of AhGβ subunits

Genomic, coding (CDS), and protein sequences of G-protein Gβ subunits from *Arachis hypogaea* L. were retrieved from the Peanut Base database (https://www.peanutbase.org/) using BLASTp, with *Arabidopsis thaliana AGB1* as query. Gene-specific primers were then designed for each gene based on the CDS sequences (Supplementary Table 4). The PCR amplified products were cloned into the pCR8 vector using the pCR™8/GW/TOPO® TA Cloning® Kit (Invitrogen, Cat. No. K250020), and the sequences were confirmed by Sanger sequencing.

To generate the phylogenetic tree of G-proteins, full-length protein sequences were obtained from sequencing results and publicly available databases (TAIR and Peanut Base). Multiple sequence alignment was performed using CLUSTALW with default parameters in MEGA X, and the aligned sequences were used to construct a maximum-likelihood phylogenetic tree with 1,000 bootstrap replicates and other default settings. The resulting tree was visualized using iTOL v7 (https://itol.embl.de/). The comparative genomic analysis was performed using TBtools II v2.330 (Chen *et al.*, 2023). Orthologous gene pairs between cultivated groundnut (*A. hypogaea*) and its wild progenitors (*A. duranensis* and *A. ipaensis*) were identified using the MCScanX toolkit in TBtools II (Chen *et al.*, 2023). Syntenic gene pairs within *Arachis hypogaea* were identified using the Multiple Collinearity Scan toolkit (MCScanX) in TBtools II (Chen *et al.*, 2023).

**Method S4: Localization studies of AhGβs**

Localization studies performed using gene-EYFP fusion constructs, full-length protein-coding regions were cloned into pEarleyGate101 vector. The AtPIP2A–mCherry construct (ABRC, Stock No. pm-rk/CD3-1007) was used as a plasma membrane marker. *Agrobacterium tumefaciens* strain GV3101 carrying the respective constructs was co-infiltrated into *Nicotiana benthamiana* leaves. Fluorescence imaging was performed using a Leica TCS SP8 spectral confocal laser scanning microscope. Nuclear staining was performed using 20 µg/ml DAPI prior to imaging. The empty vector was used as a negative control, while Arabidopsis AGB1 served as a positive control. All fluorescence images were analyzed using Fiji (ImageJ) software (Schindelin *et al.*, 2012).

**Method S5: Phenotyping and drought experiments in Arabidopsis complementation lines**

The morphological feature was analysed by growing the plants in short-day conditions (10h light/14h dark). Rosette morphology and leaf shape were assessed from two-week-old plants. Photographs were taken, and the diameter of rosette and leaf length-to-width ratio were quantified using ImageJ software. For silique morphology, seedlings were grown at 22°C under long-day conditions (16 h light/8 h dark) for 4 weeks. Mature siliques were harvested, imaged, and the silique length was measured using ImageJ. For hypocotyl analysis, seeds were surface sterilized and germinated on ½ MS medium containing 1% (w/v) sucrose and 0.8% (w/v) agar. After 2 days of stratification at 4°C, plates were incubated in complete darkness at 22°C for 3 days. Seedlings were photographed, and hypocotyl length was measured using ImageJ. To determine the effect of abscisic acid (ABA) on seed germination, seeds were surface sterilized and sown on half-strength Murashige and Skoog (½ MS) medium supplemented with 1% (w/v) sucrose and 0.8% (w/v) agar, either without ABA (control) or with 3 µM ABA. After stratification at 4°C for 2 days, plates were transferred to a growth chamber maintained at 22°C under a short-day photoperiod. Germination was scored daily for four days, using radicle emergence through the seed coat as the criterion, as observed under a stereomicroscope.

For drought-stress experiments, plants were grown at 22°C under short-day conditions (10h light/14h dark) for two weeks in a controlled growth chamber. Drought stress was imposed by withholding water for 20 days, while control plants were regularly watered to maintain uniform soil moisture. Following the stress period, plants were rehydrated, and survival rates were calculated as the percentage of recovered plants relative to the total number of treated plants.  To further assess water retention capacity, rosette leaves from 4-week-old plants were excised and immediately weighed to obtain fresh weight using a microbalance. Leaves were then placed on filter paper in a controlled environment (21-22°C, ~40% relative humidity) and weighed at 30-minute intervals for 5 hours. Water loss at each time point was expressed as a percentage of the initial fresh weight to determine the rate of dehydration.

**Method S6: Phenotyping and drought stress experiment using transgenic groundnut lines**

Phenotypic evaluation was performed using T2 plants of *AhGβ1* overexpression, *RNAi,* and WT under both well-watered and drought-stressed conditions. Plants were grown under greenhouse conditions with natural daylight variation, maintaining average day/night temperatures of 28/24°C and relative humidity levels of 60/80%. A completely randomized block design was implemented to ensure unbiased assessment of phenotypic and physiological traits in both control (well-watered) and drought-stressed. Three independent experimental blocks were used to strengthen data reliability. Drought stress was imposed at the flowering stage (40 Days after germination) by withholding water until soil moisture content (SMC) declined to below 10%, as monitored using a hygrometer, while control plants were maintained at 45-50% SMC (Supplementary figure 7F). Stress conditions were maintained for 33 days, after which plants were rehydrated to assess recovery responses. Recovery percentage was calculated as the ratio of recovered plants to the total number of plants evaluated. During the drought and recovery days, the physiological measurements, including stomatal conductance (gsw), total transpiration (E apparent), and photochemical efficiency of photosystem II under light-adapted conditions (PhiPS2), were recorded using an LI-600 portable porometer integrated with a fluorometer (LI-COR Biosciences, Lincoln, USA). Yield-associated traits were quantified for each plant, including (1) plant height, (2) total number of branches, (3) peg number, (4) pod number, (5) total seed count, (6) seed width (mm), and (7) 10-seed weight (g). Data were recorded at 60 days after germination (DAG) for the first two traits, which coincided with the final day of the drought stress period, and for all other traits at the time of harvest.

**Method S7: Transcriptome analysis**

The total RNA was extracted, and mRNA was purified using oligo-dT beads (TruSeq RNA Sample Preparation Kit, Illumina). TruSeq RNA Library Prep Kits for Illumina® (NEB, USA) were used to generate sequencing libraries based on the manufacturer’s instructions.

Libraries were sequenced using 150 bp paired-end reads on the Illumina NovaSeq 6000 platform. Raw data were pre-processed using fastp (Chen *et al.* 2018) to obtain clear reads. Filtered reads were aligned to the *Arachis hypogaea* reference genome (https://www.ncbi.nlm.nih.gov/datasets/genome/GCF_003086295.3/) using HISAT2 v2.2.1 (Kim *et al.*, 2019) with default parameters. Aligned reads used for reference-based assembly and transcript quantification with Stringtie v2.2.1 (Pertea *et al.*, 2015). Differential expression analysis was performed using the DESeq2 R-package. Differentially expressed genes selected by applying a significance threshold of p-value ≤ 0.05, with log2 fold-change cutoffs of ≥ 2 for upregulation and ≤ -2 for downregulation.

**Method S8: Construction of the transcription factor-target gene regulatory network**

Transcription factors are the major regulators of gene expression. To elucidate potential transcriptional regulatory relationships among the differentially expressed genes (DEGs), a comprehensive TF-target prediction pipeline was implemented. The promoter sequences (2.5 kb upstream of the transcription start site) corresponding to DEGs were retrieved using GPXplore tool (Godara et al., 2026). Transcription factor binding site (TFBS) prediction was performed using the Regulation Prediction module of PlantRegMap (https://plantregmap.gaolab.org/regulation_prediction.php). Due to limited TF-target annotations and incomplete regulatory models for *Arachis hypogaea*, motif libraries from its two diploid progenitor species (*Arachis ipaensis* and *Arachis duranensis*) were used. In addition, *Arabidopsis thaliana* TFs were included as a supporting dataset to enhance the sensitivity of motif detection. PlantRegMap identifies TF-target interactions by scanning promoter sequences with high-confidence TF binding motifs using the FIMO algorithm, and an interaction is assigned when at least one statistically significant (pvalue <1e-5) binding site is detected within the promoter. Predicted regulation tables were curated to ensure unique TF-target representation. For example, if a TF contained multiple predicted binding sites within the same promoter, only the most proximal site (closest to the transcription start site) was retained to avoid duplications of regulatory edges.

To identify TF orthologs of *A. hypogaea*, TF identifiers from *A. ipaensis, A. duranensis*, and *A. thaliana* were mapped to their corresponding orthologs in the *A. hypogaea* genome. Ortholog mapping was performed using the Best ID Conversion tool available in TBtools II. The predicted TF orthologs were then cross-referenced with the DEG list to identify TFs that might be regulated downstream of *AhGβ1*. The TF orthologs that overlapped with the DEG set were subsequently used to filter the predicted TF-target interactions from three species, ensuring that only regulatory relationships involving DEGs were retained. TF-target pairs from all three species were subsequently merged to construct a unified regulatory network for *A. hypogaea*. Redundant interactions arising from multiple species predicting the same regulation were collapsed into unique TF-target pairs. This merged, non-redundant network was considered the final predicted transcriptional regulatory network of A. hypogaea. The predicted TF-target network was further examined to understand the overall regulatory patterns within the network. To further increase confidence in the predicted regulations, the TF-target network was integrated with the gene co-expression network derived from Pearson correlation analysis (|R| >0.7 and adj_pvalue<0.05). Overlapping regulatory edges were identified, and the combined TF-coexpression network was visualized and explored in Cytoscape to highlight consistent regulatory signals supported by both promoter motif analysis and expression-based co-regulation. The regulatory modules were further analyzed using GO-DIRECT and KEGG pathways in the DAVID Bioinformatics tool (https://davidbioinformatics.nih.gov/tools.jsp).

**Method S9: Untargeted metabolome analysis**

Chromatographic separation was performed on a C18 Hypersil Gold column using a standardized gradient elution program. Both positive and negative electrospray ionization (ESI) modes were performed for all samples with pooled quality control (QC) and blank. Raw LCMS data files were converted to mzML format and processed using the XCMS (v4.8) R-package. The processing includes peak alignment, retention time correction, peak grouping, gap filling, and blank correction. Features with >50% missing values and QC coefficient of variation >30% were removed prior to downstream statistical analysis.

**Method S10: Total flavonoid content and $H_2O_2$ quantification**

Total flavonoid content (TFC) was quantified using the aluminum chloride colorimetric method (Baba and Malik, 2015). One hundred mg of tissue was used for extraction in 500 µl of methanol. 10 µl of crude extract was mixed with 90 µl of methanol and 400 µl of distilled water. To this, 30 µl of 5% $NaNO_2$, and 30µl of 10% $AlCl_3$ solutions were added, and the mixture was incubated for 6 min. Later, 200 µl of 1 M NaOH solution was added to make up the volume to 1 ml with distilled water. After 15 min incubation, absorbance was recorded at 510 nm. The total flavonoid content was measured using a standard curve of quercetin (Qt) and expressed in µgQt/mg dry weight.

$H_2O_2$ quantification was carried out with the ferrous ammonium sulfate/xylenol orange (FOX) method (Sakil *et al.*, 2023). Fresh leaves (100 mg) were ground using liquid nitrogen, followed by homogenized in 1 ml of 25 mM $H_2SO_4$ and centrifuged at 15,000×g for 5 minutes. Approximately 100 µl supernatant was mixed with 1 ml of FOX reagent, prepared with 150 µM ammonium ferrous sulfate, 150 µM xylenol orange, and 100 mM sorbitol in 25 mM $H_2SO_4$. After incubation in the dark for 20 minutes, the absorbance was recorded at 560 nm. The $H_2O_2$

concentration was calculated using a standard curve and expressed as μmol $H_2O_2$ per gram of fresh weight.

### Method S11: Lignin and cellulose staining

Leaf petioles were collected and fixed in 4% paraformaldehyde solution, and 100 μm sections were prepared using a vibrotome (VT1000 S; Leica). Lignin staining performed using Wiesner reagent (Johansen, 1940) and images were acquired under compound microscope. Cellulose staining performed using Calcofluor White dye (Bidhendi *et al.*, 2020) and Leica TCS SP8 spectral confocal laser scanning microscope used for imaging with 405 nm excitation laser and emission collected at 430-460 nm. Mean fluorescence intensity measured in Fiji/ImageJ (Schindelin *et al.*, 2012).

### Method S12: DPBA staining and confocal imaging of flavonol-associated fluorescence

Leaf samples were collected 5 days after foliar treatment with quercetin, rutin, and DMSO and used for flavonol visualization using diphenylboric acid 2-aminoethyl ester (DPBA) staining using the method described earlier (Nguyen, 2020). Briefly, epidermal peels were prepared from groundnut leaves and stained with 0.25% (w/v) diphenylboric acid 2-aminoethyl ester (DPBA). Samples were vacuum infiltrated with DPBA solution in the dark at room temperature, washed with PBS, and mounted in 20% glycerol. Unstained samples were used to assess background autofluorescence. DPBA-stained epidermal peels were imaged using a Leica TCS SP8 confocal microscope under identical acquisition settings for all treatments. DPBA-associated fluorescence was recorded at 577-620 nm following 488-nm excitation. Chlorophyll fluorescence was detected at 650-700 nm following 561-nm excitation. Fluorescence intensity was quantified from the original image data using Image J, corrected by subtracting the corresponding unstained background, and normalized to the number of stomata in each field.
